# Contemporary clinical isolates of *Staphylococcus aureus* from pediatric osteomyelitis patients display unique characteristics in a mouse model of hematogenous osteomyelitis

**DOI:** 10.1101/2021.03.22.436444

**Authors:** Philip M Roper, Kara R Eichelberger, Linda Cox, Luke O’Connor, Christine Shao, Caleb A Ford, Stephanie A Fritz, James E Cassat, Deborah J Veis

**Affiliations:** Division of Bone & Mineral Diseases, Musculoskeletal Research Center, Washington University School of Medicine, Saint Louis, MO, USA; Department of Pathology and Immunology, Washington University School of Medicine, Saint Louis, MO, USA; Department of Pathology, Microbiology, and Immunology, Vanderbilt University Medical Center, Nashville TN, USA; Department of Pediatrics, Division of Pediatric Infectious Diseases, Vanderbilt University Medical Center, Nashville, TN, USA; Department of Biomedical Engineering, Vanderbilt University, Nashville, TN, USA; Vanderbilt Institute for Infection, Immunology and Inflammation (VI4), Vanderbilt University Medical Center, Nashville, TN, USA; Vanderbilt Center for Bone Biology, Vanderbilt University Medical Center, Nashville, TN, USA; Shriners Hospitals for Children, Saint Louis, MO, USA; Department of Pediatrics, Division of Pediatric Infectious Diseases, Washington University School of Medicine, Saint Louis, MO, USA

## Abstract

Osteomyelitis can result from the direct inoculation of pathogens into bone during injury or surgery, or from spread via the bloodstream, a condition called hematogenous osteomyelitis (HOM). HOM disproportionally affects children, and more than half of cases are caused by *Staphylococcus (S.) aureus*. Laboratory models of osteomyelitis mostly utilize direct injection of bacteria into the bone or the implantation of foreign material, and therefore do not directly interrogate the pathogenesis of pediatric hematogenous osteomyelitis. In this study, we inoculated mice intravenously and characterized resultant musculoskeletal infections using two strains isolated from adults (USA300-LAC and NRS384) and five new methicillin-resistant *S. aureus* isolates from pediatric osteomyelitis patients. All strains were capable of creating stable infections over five weeks, although the incidence varied. Micro-computed tomography (microCT) analysis demonstrated decreases in trabecular bone volume fraction but little effect on bone cortices. Histologic assessment revealed differences in the precise focus of musculoskeletal infection, with varying mixtures of bone-centered osteomyelitis and joint-centered septic arthritis. Whole genome sequencing of three new isolates demonstrated distinct strains, two within the USA300 lineage and one USA100 isolate. Interestingly, the USA100 strain showed a distinct predilection for septic arthritis, compared to the USA300 strains, including NRS384 and LAC, which more frequently led to osteomyelitis or mixed bone and joint infections. Collectively, these data outline the feasibility of using pediatric osteomyelitis clinical isolates to study the pathogenesis of HOM in murine models and lay the groundwork for future studies investigating strain-dependent differences in musculoskeletal infection.

**Importance:** The inflammation of bone tissue is called osteomyelitis, and more than half of cases are caused by an infection with the bacterium *Staphylococcus aureus*. In children, the most common route of infection is hematogenous, wherein bacteria seed the bone from the bloodstream without another known site of infection. Although these infections pose a significant health problem, they are understudied in the laboratory because of a dearth of robust animal models. In this study, we utilized several previously uncharacterized clinical isolates of *S. aureus* derived from children with bone infections to generate reproducible and stable musculoskeletal infection in mice with many features seen in human osteomyelitis, making them a valuable resource for future mechanistic and therapeutic studies.

## Introduction

Infectious osteomyelitis is commonly bacterial in origin, with *Staphylococcus (S.) aureus* the most frequent pathogen identified (1). Hematogenous osteomyelitis (HOM), initiated via seeding of bacteria through the bloodstream, predominantly affects children, with 85% of cases occurring in children under 17 years of age (1–4). HOM in children is not rare, with reported incidence between 1 in 1000 and 1 in 20,000 (5–7), but it is notoriously difficult to treat, often requiring prolonged antibiotics and surgical debridement. The recurrence rate may be as high as 9% (7–10), and this problematic event can extend into adulthood (11). Typically, HOM involves the metaphysis of long bones, predominantly the distal femur or proximal tibia (12). A main concern with pediatric HOM is the development of chronic infection. This can cause pathological bone changes leading to fractures, dislocations, and devitalized bone segments known as sequestra (12). HOM infections can also spread to an adjacent joint, causing septic arthritis (SA), or systemically, leading to potentially lethal sepsis (12, 13). Additionally, antimicrobial resistant strains, particularly methicillin-resistant (MRSA), are highly prevalent in both healthcare and community settings (14).

Despite significant clinical challenges, studies of the specific mechanisms underlying HOM development and progression have been relatively limited. This is partly because most animal models recapitulate cases of bone infection related to orthopedic surgery and involve manipulations of the bone including creation of cortical defects or placement of hardware into the bone (15–20). One study utilized the hematogenous route but relied upon the implantation of metal into the femur before the injection of bacteria (19). Although HOM has been modeled in mice without prior manipulation (21–24), characterization of bone changes and comparisons among strains have been limited.

In addition to two previously established community-acquired USA300 strains isolated from adults, (16, 25, 26), we utilized several new clinical isolates of *S. aureus* from pediatric HOM patients. To mimic the bone microenvironment in children, the population in which HOM is most prevalent, we intravenously inoculated actively growing mice. Generation of stably bioluminescent bacteria allowed for real-time, longitudinal assessment of infection localization and progression *in vivo* while preserving the bone for downstream analysis by micro-computed tomography (microCT) and histology. We compared measures of pathogenicity between these isolates, including incidence of hindlimb infection, propensity towards developing bone-centered HOM or joint-centered septic arthritis (SA), subsequent bone loss, cytotoxicity toward host cells, and intracellular proliferation. We also performed genomic sequence analysis of 3 of these isolates, discovering significant genetic differences amongst the clinical isolates. Taken together, these studies provide a robust platform for the analysis of host-pathogen interactions in HOM.

## Results

### Modeling hematogenous osteomyelitis using NRS384 and LAC

We first established HOM with two USA300 strains of *S. aureus* previously made spontaneously bioluminescent by us and another group, via stable integration of a plasmid bearing the *lux* operon (27, 28) allowing tracking of infection over time *in vivo*. We injected 2.5×10^6^ colony forming units (CFU) of NRS384 or LAC in PBS into the tail vein of wild type C57BL6-albino mice at 6 weeks of age, a period of rapid growth. With this inoculum, we did not observe any instances of lethal sepsis or other complications requiring euthanasia prior to 4 weeks post infection with either strain (Fig S1). Bioluminescence imaging (BLI) was performed weekly for 5 weeks. With both USA300 strains, distinct foci of signal were frequently seen in hindlimbs as well as bladder, likely indicative of bacteria being shed from the kidneys (Fig. 1A,B). Foci of signal were also observed in the rostrum and occasionally the forelimb. Based on this distribution of bone-centric signal, regions of interest were drawn over the hindlimbs, also referred to as legs, (as in Fig. S2), allowing quantitation over time (Fig. 1C,D). We found no appreciable differences between signal intensities from two cohorts of mice injected on different days (Trial 1 and Trial 2), and the signal values stabilized around the 2-week timepoint, suggesting establishment of chronic infection. Because of this BLI signal stabilization, the Trial 2 LAC-injected mice were harvested at 2 weeks. Next, we used all the BLI values acquired to devise a BLI positivity threshold to stratify actively infected BLI positive (+) legs from BLI negative (-) legs for later analyses. This threshold was set at 70,000 photons/sec, because legs under this value consistently had no foci of BLI signal over the 5-week course of infection, excluding “bleed” from strong bladder signals (as in Fig. S2B, right leg), while the baseline signal for autoluminescence was around 35,000 photons/sec. Further, no legs that reached this threshold subsequently fell below it. Interestingly, we observed that the cohorts injected with NRS384 had a markedly higher percentage of mice that developed a hindlimb infection detectable by BLI compared to LAC-injected cohorts (80% vs. 35% cumulative incidence).

**Figure 1.**
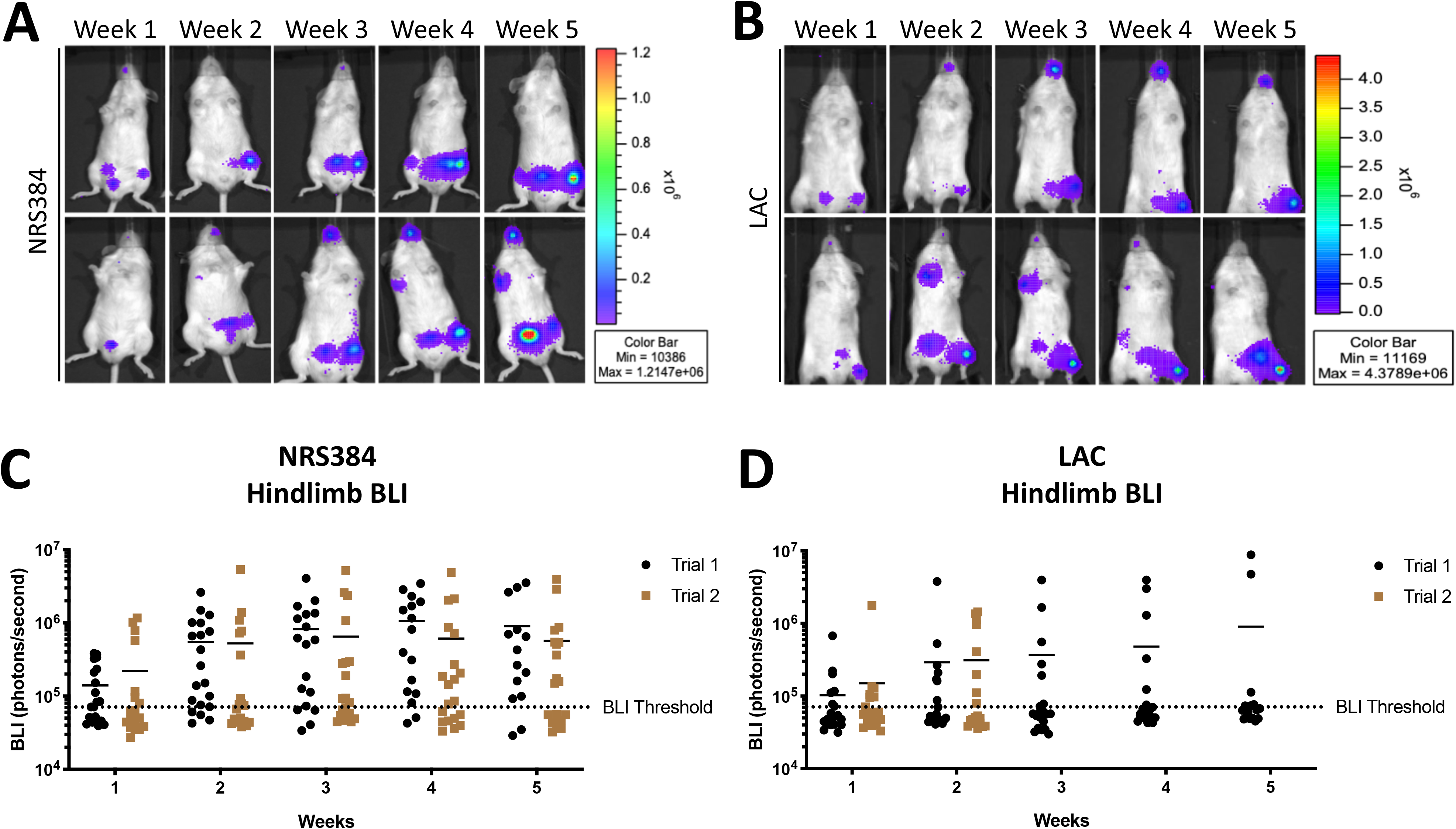
Longitudinal tracking of NRS384 and LAC hematogenous osteomyelitis. *In vivo* bioluminescent imaging (BLI) of two representative mice (top and bottom rows), performed weekly for 5 weeks, after injection with NRS384 (A) or LAC (B) strains of *S. aureus*. The quantification of NRS384 (C) and LAC (D) BLI signal measured in each hind limb (2 data points per mouse) for two separate trials is depicted over the 5 weeks post-injection (n=10 mice per trial; BLI Threshold is set at 70,000 p/sec/cm^2^/sr). LAC Trial 2 purposely terminated at 2 weeks post-injection.

Since most BLI signal for NRS384-injected mice appeared to be proximal to the knee joint, femurs were assessed by microCT. Reconstructions indicated that the bone disruption is predominantly localized to the trabecular compartment (Fig. S3A). Quantification of the trabecular bone volume fraction (ratio of bone volume to total volume, or BV/TV) in the femur demonstrated significant decreases from both BLI- and BLI+ legs, compared to those from non-infected control mice (Fig. 2A). While no statistical difference was found between BLI- and BLI+ legs of these NRS384-infected mice, extreme destruction of trabecular bone (BV/TV below 0.1) was only seen in BLI+ legs. Interestingly, BLI signal from LAC-injected mice appeared more distal than in NRS384-injected mice, so we measured the trabecular bone volumes from both femurs and tibias. The femoral BV/TV in LAC-injected mice was lower than control mice, but there was no difference in BV/TV between BLI- and BLI+ legs, nor between those harvested at 2 or 5 weeks (Fig. 2B, S3B). However, the trabecular bone volumes in tibias of BLI+ legs were decreased compared to tibias from those that were BLI-, reaching statistical significance at 5-weeks post-injection (Fig. 2C, S3B). Overall, we were able to induce consistent hindlimb HOM with both the NRS384 and LAC strains using our intravenous injection model.

**Figure 2.**
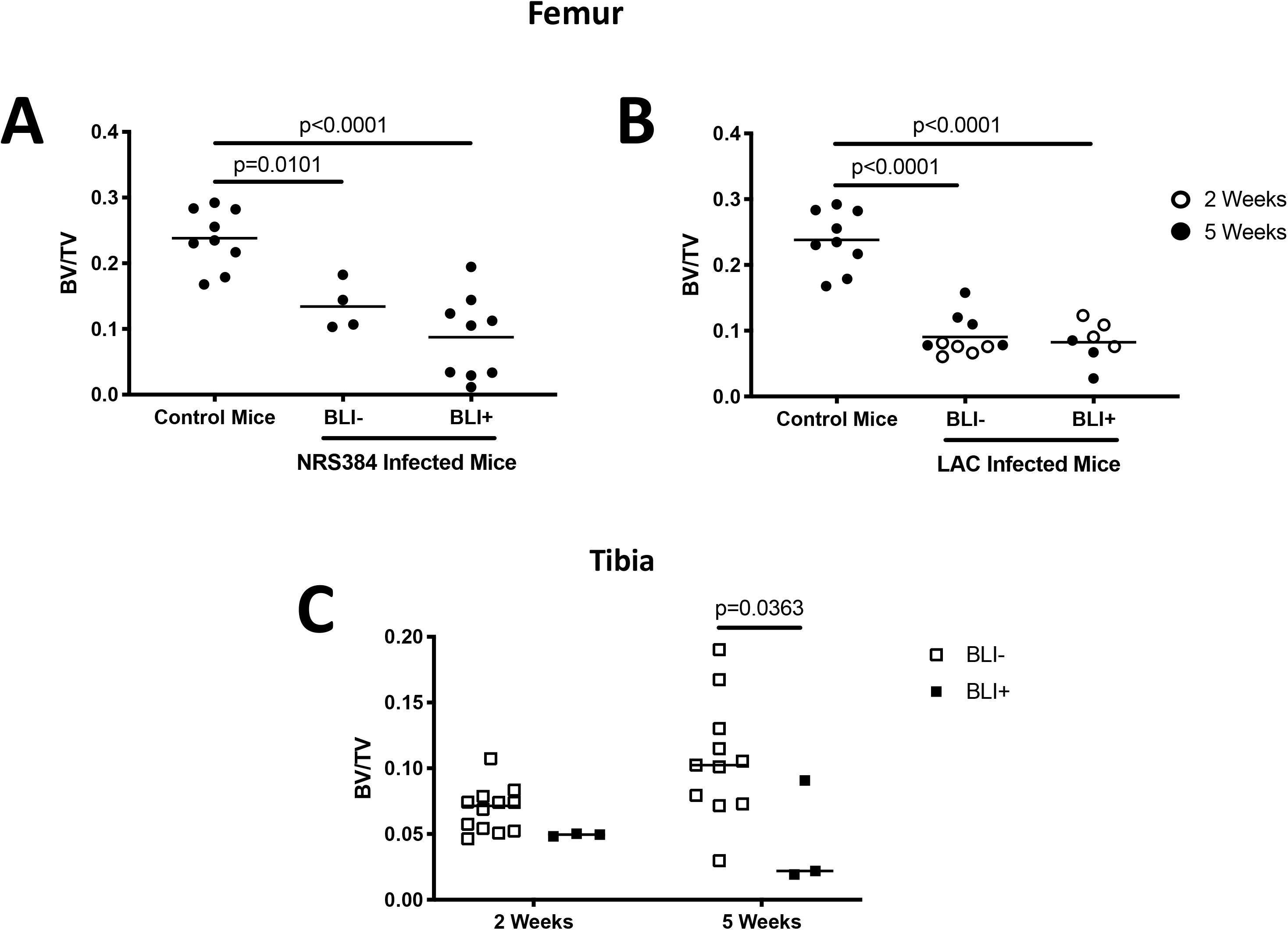
Trabecular bone volume fractions from NRS384- and LAC-injected mice. Trabecular bone volume/total volume (BV/TV) was measured by micro-computed tomography (microCT) in femurs from uninfected control mice and NRS384 (A) and LAC (B) injected mice. The control group (age, sex and strain-matched) is the same in A and B. (C) Tibial trabecular BV/TV from LAC injected mice was measured at 2- or 5-weeks post-injection. BLI- or BLI+ designation is based on the threshold in Figure 1. P values were calculated by one-way ANOVA with Tukey’s Multiple Comparisons post-hoc test.

### Characterizing HOM caused by *S. aureus* clinical isolates from children with OM

To expand the clinical significance of this model to pediatric HOM, we acquired five isolates of methicillin-resistant *S. aureus* (MRSA) from pediatric patients with HOM (Table 1). In order to employ these isolates for BLI imaging during HOM in mice, we utilized the same stable chromosomal integration system for the *lux* operon used previously for NRS384 (27). To confirm that the bioluminescence of each isolate corresponds to an equivalent bacterial burden, we measured the BLI signal from *in vitro* serial dilutions of each isolate and correlated that with CFUs enumerated from the same dilutions (Fig. S4). In liquid culture, growth of TI2 and TI5 were minimally delayed (Fig. S5). Tail vein injection of 2.5×10^6^ CFU of all 5 isolates led to reproducible infections with localization over hindlimbs (Fig 3A, S6A,B) and limited lethality (Fig. S6C), comparable to that of NRS384 and LAC (Fig. S1). Interestingly, we discovered differences between the isolates in the rate at which they led to hindlimb infections in these mice, using our previously established BLI positivity threshold (Fig. 3B). Two of the isolates, TI1 and TI3 had a very high incidence of infection in at least one hindlimb, 72% and 89%, respectively, comparable to NRS384. However, TI2 had a much lower rate of infection, 25%, over the 5-week time course, similar to the incidence with LAC. The other two strains, TI4 and TI5 had infection rates between these extremes, 46% and 67%, respectively. For subsequent analyses we focused primarily on TI3 and TI1, with the highest incidence of hindlimb infection, as well as TI2, with the lowest.

**Figure 3.**
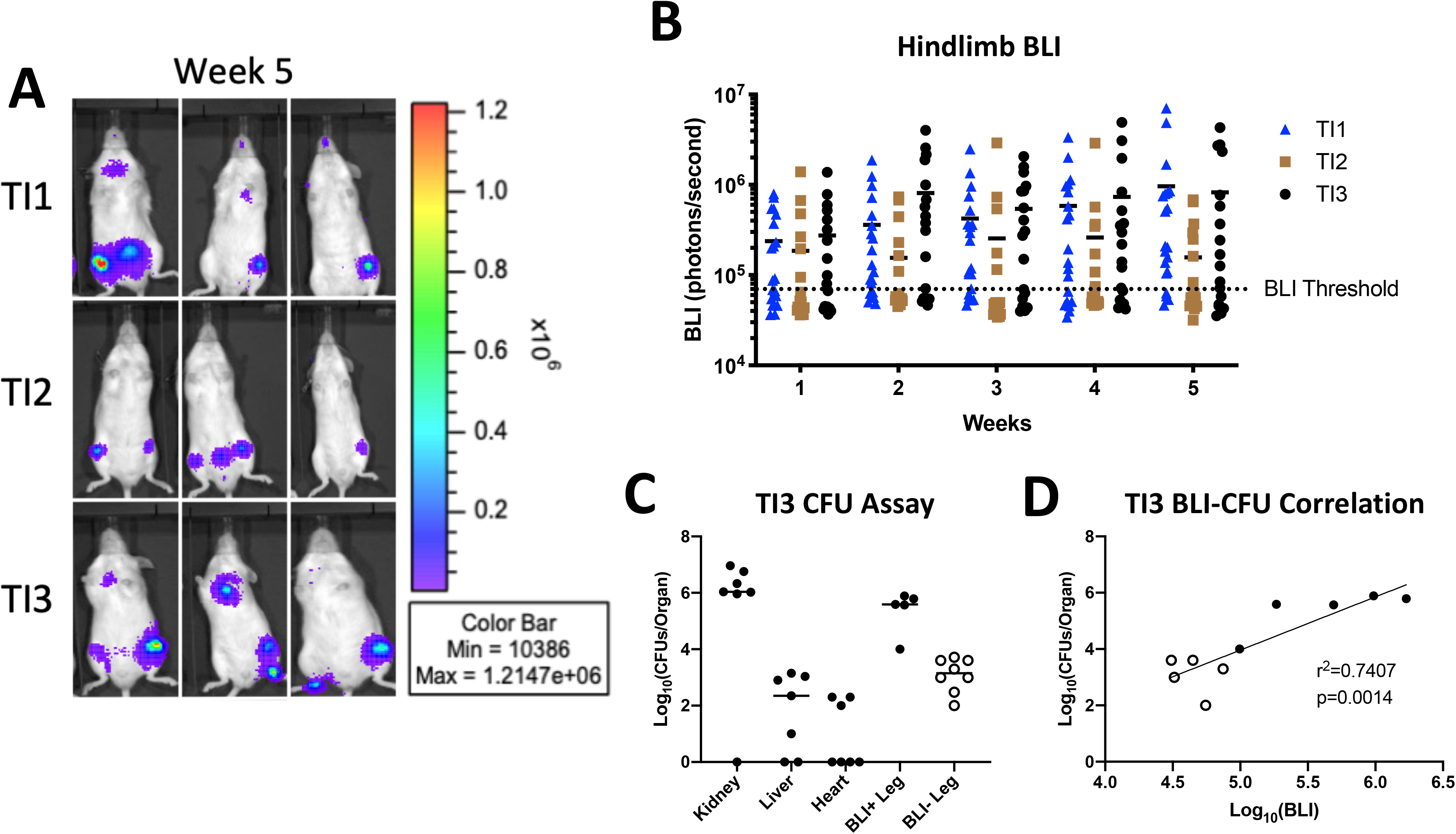
Characterization of hematogenous osteomyelitis using clinical isolates of *S. aureus*. (A) *In vivo* bioluminescent imaging (BLI) of three representative mice for each isolate at 5 weeks post-injection. (B) The quantification of BLI signal measured in the hind limb of each injected mouse depicted over the 5 weeks post-injection (n=20 per isolate; BLI Threshold 70,000 photons/second). (C) Enumerated bacterial burdens from designated organs, as measured by colony forming unit (CFU) assay at 3 weeks post-injection. BLI+ and BLI- Legs are the counts from femurs from legs with BLI signal higher or lower than 70,000 photons/sec, respectively (n=7 mice from a separate cohort from (A) and (B) injected with TI3, each dot represents one leg). (D) Correlation of the bacterial burden (CFU) from (C) with the BLI signal of the corresponding leg. Open and closed dots correspond to the BLI+/- designation in (C).

**Table 1.**
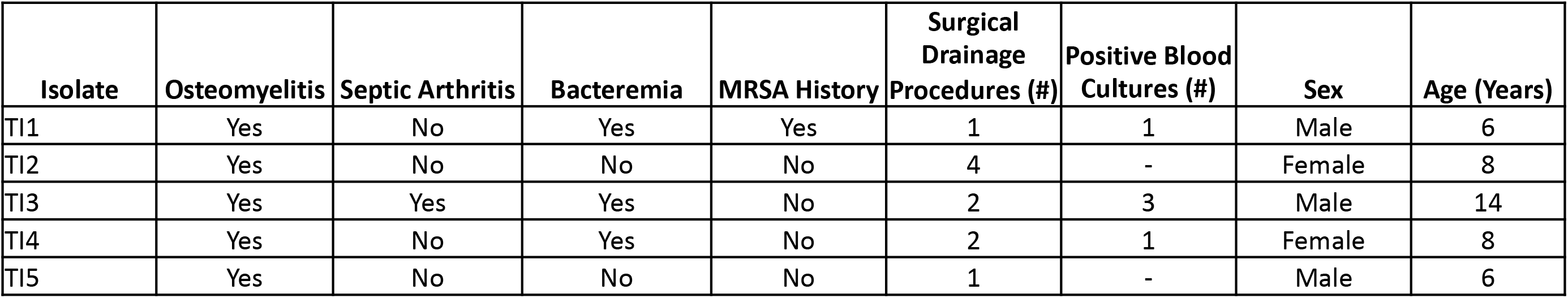
Source information for *S. aureus* clinical isolates. Each of the five clinical isolates used throughout these studies were obtained from white, non-Hispanic juvenile patients at Saint Louis Children’s Hospital, St. Louis, MO.

| Isolate | Osteomyelitis | Septic Arthritis | Bacteremia | MRSA History | Surgical Drainage Procedures (#) | Positive Blood Cultures (#) | Sex | Age (Years) |
| --- | --- | --- | --- | --- | --- | --- | --- | --- |
| TI1 | Yes | No | Yes | Yes | 1 | 1 | Male | 6 |
| TI2 | Yes | No | No | No | 4 | - | Female | 8 |
| TI3 | Yes | Yes | Yes | No | 2 | 3 | Male | 14 |
| TI4 | Yes | No | Yes | No | 2 | 1 | Female | 8 |
| TI5 | Yes | No | No | No | 1 | - | Male | 6 |

To further validate BLI as a proxy for bacterial load *in vivo*, we measured CFUs obtained from homogenized femurs, along with the heart, liver, and kidneys, from mice injected with TI3 two weeks prior (Fig. 3C). High levels (∼5×10^6^ CFU/organ) of bacteria were found in the kidneys and hind limb bones from BLI+ legs, as expected, and the correlation between CFUs and the corresponding BLI signal in hindlimbs was strong (Fig. 3D).

We next compared the trabecular BV/TV from mice injected with these three clinical isolates, focusing on femurs because BLI signals were primarily proximal to the knee. As with the NRS384-injected mice, we found a significant decrease in all the bone volumes of femurs from mice injected with each clinical isolate compared to femurs of non-injected mice. However, with these mice we also found a consistent decrease in the bone volume fractions of femurs from BLI+ legs as compared to BLI- legs (Fig. 4A). Finally, we pooled the data from the BLI+ and BLI- legs of all three clinical isolates and found a weak but statistically significant correlation between the BLI signal and the BV/TV from the corresponding femur (Fig. 4B). These data further validated our utilization of BLI to track infection incidence and location.

**Figure 4.**
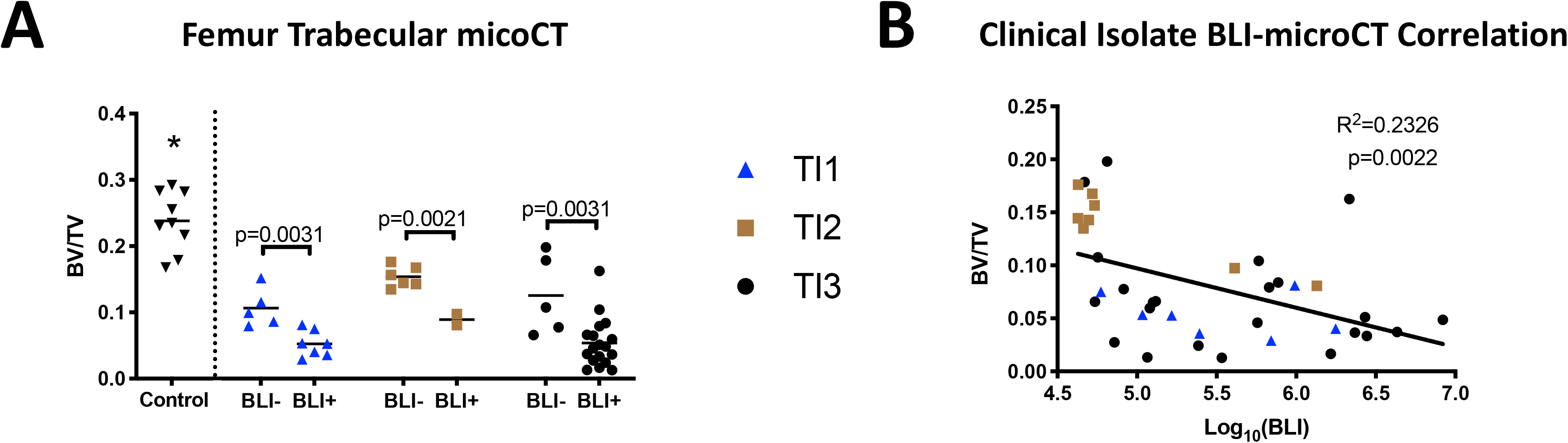
Reduced bone volumes are associated with local bacterial burden with HOM induced by clinical isolates. (A) Trabecular BV/TV measured by microCT of femurs from non-injected control mice (same cohort as in Figure 2) and mice injected with either TI1, TI2, or TI3. Classification as BLI- or BLI+ legs was determined by the threshold in Figure 3. *p≤0.0014, Control vs all other groups as calculated by one-way ANOVA with Tukey’s multiple comparison *posthoc* test. P values between BLI- and BLI+ BV/TV within each strain were calculated by Student’s t-test. (B) Correlation of the BV/TV data from (A) with the BLI signal from the corresponding leg. Dot coding corresponds to the coding in (A).

### Osteomyelitis and septic arthritis result from hematogenous inoculation with *S. aureus strains*

We next examined hindlimbs by standard histology 5 weeks after infection, finding varying combinations of bone and joint involvement. Figure 5 depicts representative histological images of three separate samples of the distal femur and knee joint (one per column) from infections with NRS384 and LAC and three pediatric isolates (TI1, TI2, and TI3). Some samples (Fig. 5D, H, M and N) appear to have the infections localized entirely within the bone and medullary cavity, indicative of OM. Other samples (Fig. 5B, I, K, L) show clear infection and inflammation within the joint space (Black Arrows) while the femur and tibia remain normal, demonstrating septic arthritis (SA). Finally, there are samples (Fig. 5A, C, E, F, G, O) where the infection is present both in the bone and joint (OM+SA). However, in these latter samples, some infections seem to extend into the joint from severe bone involvement (Fig. 5A, C, F; black arrowheads), while others seem to represent a joint-centered infection focally extending into bone (Fig. 5J; black arrowheads). In all cases of OM infections, there is significant loss of trabecular bone, and in extreme cases there is disruption of the growth plate as well (Fig. 5C, D, E, J, M, O; #). We did not observe extension through the cortical bone to the periosteal surface in affected femurs. The various patterns of bone and joint involvement for each strain were then enumerated (Table 2). NRS384 and TI1 produced mixtures of bone-centered OM and joint-centered SA infections. However, LAC and TI3 predominantly produced OM, with no case of SA that was not coincidental with OM. TI2 predominantly produced SA, with only one case of OM that was not associated with SA. Interestingly, LAC produced OM lesions in the tibias but not femurs of 6 out of 7 mice, as opposed to all other strains that consistently produced OM in the femurs.

**Figure 5.**
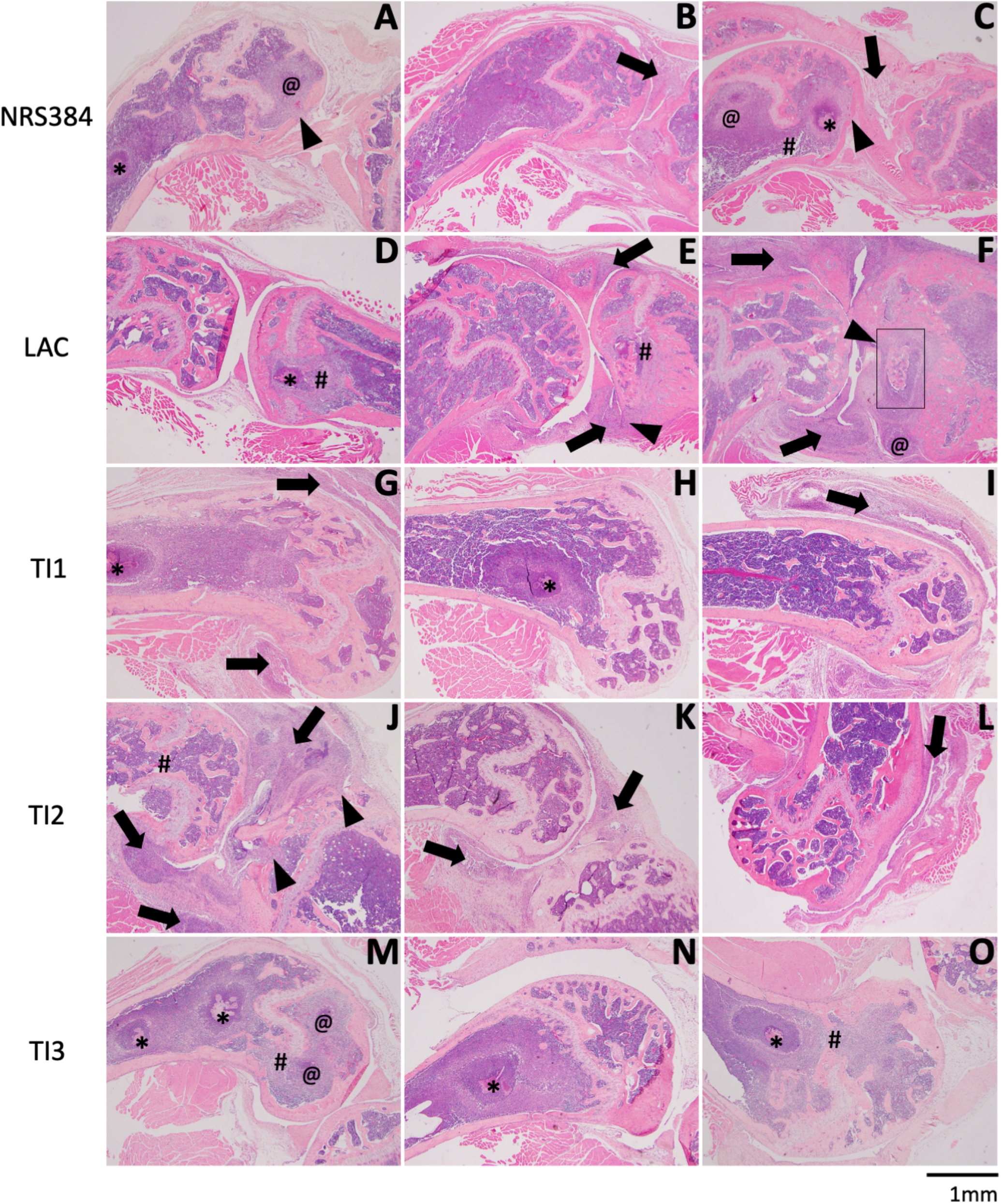
Histological changes to hind limbs with HOM. Each row depicts representative histology of the distal femur and knee joint, with or without proximal tibia, from three separate samples of mice injected with NRS384 (A-C), LAC (D-F), TI1 (G-I), TI2 (J-L), or TI3 (M-O). In panels with both bones, femur is on the left and tibia on the right. Symbols denote specific histological characteristics: *, abscess with visible bacterial colonies; @, abscess without visible bacterial colonies; # growth plate disruption; black arrows, inflammation within the joint space and/or synovium; black arrowheads, infection focally extending from joint into bone. Box in (F) depicts area of higher magnification in Fig. 6C.

**Table 2.** Infection localization in BLI+ limbs as assessed histologically. Fractions listed as (number of samples that exhibit the indicated histologic characteristic)/(total number of samples examined). Characteristics were determined as follows: Osteomyelitis (OM) if abscesses and/or inflammation was present within the bone tissue or medullary cavity; Septic Arthritis (SA) if abscesses and/or inflammation was present in the joint space; OM+SA if there was both OM and SA present; and No Changes if no overt histological changes were seen in either the bone or the joint, on at least 2-3 sections per sample.

| **Strain** | **Osteomyelitis (OM)** | **Septic Arthritis (SA)** | **OM+SA** | **No changes** |
| --- | --- | --- | --- | --- |
| NRS384 | 5/9 | 1/9 | 1/9 | 2/9 |
| LAC | 3/7 | 0/7 | 4/7 | 0/7 |
| TI1 | 4/12 | 2/12 | 3/12 | 3/12 |
| TI2 | 1/8 | 3/8 | 3/8 | 1/8 |
| TI3 | 12/18 | 0/18 | 5/18 | 1/18 |

Focusing on OM lesions, higher magnification images demonstrate characteristic abscesses with neutrophil infiltration and large macrophages surrounding the lesions (Fig 6A, B: black arrowheads), some with central bacterial colonies (Fig. 6B; *). We observed only one example of a sequestrum, in a tibia from a LAC infection (Fig. 5F; black box). In the higher magnification image, collections of cocci can be seen on the bone surface (Fig, 6C; black arrows), as well as empty lacunae. Since we saw dramatic bone loss, we looked for osteoclasts by staining for tartrate-resistant acid phosphatase (TRAP, an osteoclast marker), finding that many bone surfaces close to abscesses were covered by these resorptive cells (Fig. 6D-I).

**Figure 6.**
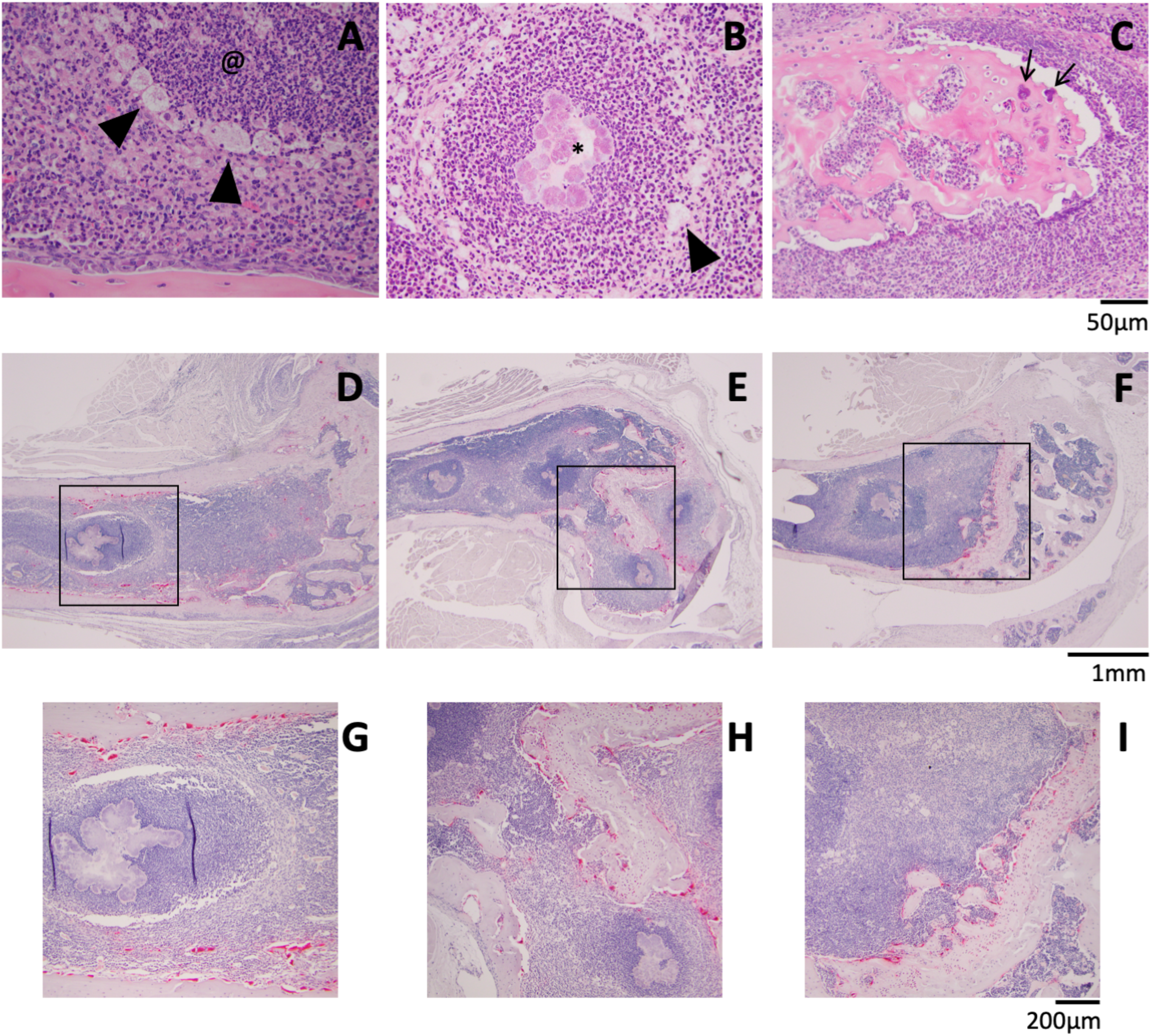
Detailed histological characteristics of HOM. Abscesses without (A; @) and with (B;*) visible bacterial colonies were surrounded by enlarged macrophages (Black Arrowheads). (C) Magnified image of sequestrum from Fig 5F showing bacteria (black arrows). (D-I) TRAP staining (red) for osteoclast activity. (G), (H), and (I) depict the higher magnification of the black boxes indicated in (D), (E), and (F), respectively. Scale bars are presented for each row of images.

While it was not uncommon to find damage to the tibial epiphysis in cases with SA, involvement of the tibial metaphysis or diaphysis was rare in all strains used except for LAC. One TI3-injected mouse had abscesses with bacterial colonies throughout the diaphyseal marrow space and a focus of periosteal reaction surrounding an abscess that appears to be outside of the normal cortical shell (Fig. S7A). In addition to displaying markedly more tibial involvement than all the other strains, LAC-injected mice also had two cases of inflammation with reactive bone formation on the periosteal surface (Fig. S7B-D). In one case, the metaphysis of a tibia from a TI2-injected mouse was fibrotic, with a rim of new bone formation, just distal to the growth plate (Fig. S7E,F). While not prevalent, these instances of new bone formation provide further evidence that our murine HOM shares many features of the human disorder. Altogether, this histology reveals significant pathologic differences of the OM infections between the clinical isolates.

### Varying effects of clinical isolates on osteoblasts and osteoclasts

Next, we investigated whether these different strains exhibited virulence differences using our osteoclast intracellular proliferation (29) and osteoblast cytotoxicity assays (16). First, osteoclasts differentiated for 3 days in culture were separately infected with each strain for 30 minutes, then all uninternalized bacteria were killed by the addition of antibiotic to the culture for 1 hour. Osteoclasts were lysed immediately thereafter (1.5 hours post infection), indicative of internalization, or cultured until 18-hours post-infection, to allow proliferation, and CFU were enumerated. TI2 produced a significantly lower increase in the CFU between 1.5 to 18 hours, as compared to all other strains (Fig. 7A). Input CFU and CFU measured at 1.5 hours after infection were similar among all strains (Fig. S8A and B), indicating the observed difference in TI2 CFU at 18 hours was related to decreased intracellular proliferation within osteoclasts and not internalization. Next, cytotoxicity was assessed by adding concentrated bacterial supernatants to cultured MC3T3 osteoblastic cells for 22 hours (Fig. 7B). While all other strains showed complete cytotoxicity with the addition of 10% supernatant, TI2 supernatant had no effect on osteoblast cell survival. LAC and TI3 were the most cytotoxic, with complete killing at 5% supernatant. Thus, the lower incidence of HOM with TI2, but not LAC, was associated with decreased direct effects on bone cells. These data, in addition to our histology, further characterize TI2 as functionally distinct from the other isolates.

**Figure 7.**
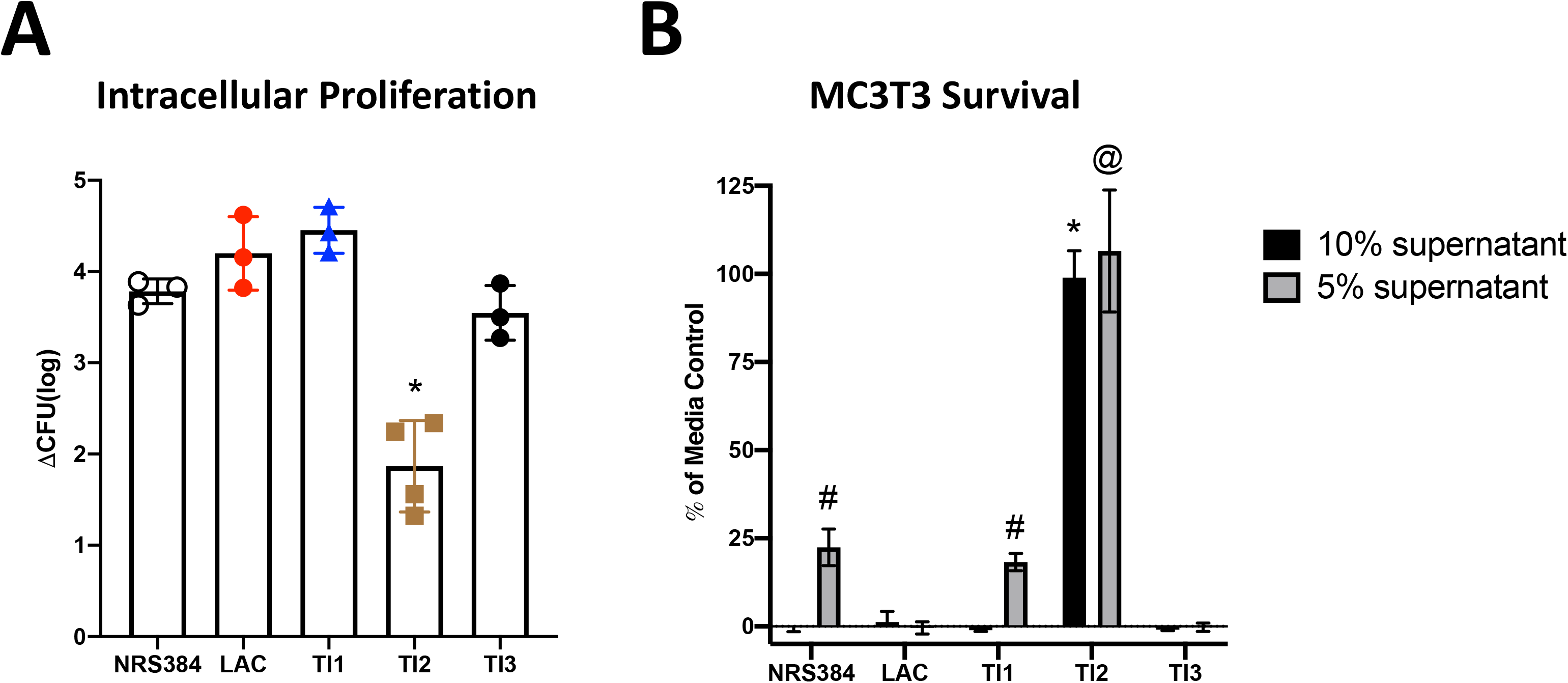
Effects of *S. aureus* strains on bone cells *in vitro*. (A) The intracellular proliferation within cultured osteoclasts measured by change in CFU between 1.5 and 18 hours post infection. *p≤0.0015 compared to all other strains as measured by one-way ANOVA with Tukey’s post-hoc test. (B) MC3T3 survival after 22 hours of exposure to media with either 5% or 10% concentrated *S. aureus* concentrated culture supernatant, relative to a media-only control. Bars represent the mean and error bars are SD. N=10; # and @p<0.0001 as compared to all other 5% supernatants, *p<0.0001 as compared to all other 10% supernatants, by two-way ANOVA with Tukey’s post-hoc test.

### Genetic sequence typing of the clinical isolates

In order to begin investigating the inherent genetic differences underlying the observed differences in pathogenicity, the genomes of three clinical isolates (TI1, TI2, and TI3) were sequenced and multilocus sequence typing (MLST) was performed as previously described (30). This analysis showed that TI2 belongs to clonal complex (CC) 5, indicative of a USA100 clonal lineage, while TI1 and TI3 are both CC ID8, consistent with a USA300 clonal lineage (Fig. 8A). When each isolate was compared with its respective clonal lineage reference strain (FPR3757 for USA300 and N315 for USA100), TI2 had more than 3 times the number of total polymorphisms, as well as genes containing at least one polymorphism, detected than TI1 and TI3 (Fig. 8B and C). Next, 173 virulence-associated genes within each isolate were grouped by function and examined for the presence of polymorphisms, as compared to their respective reference strain (31). TI2 shows more polymorphisms in genes related to adhesins and immune evasion than Tl1 or TI3 (Fig. 8D and E; S9, red squares). However, polymorphisms in the other categories were relatively infrequent in all pediatric strains.

**Figure 8.**
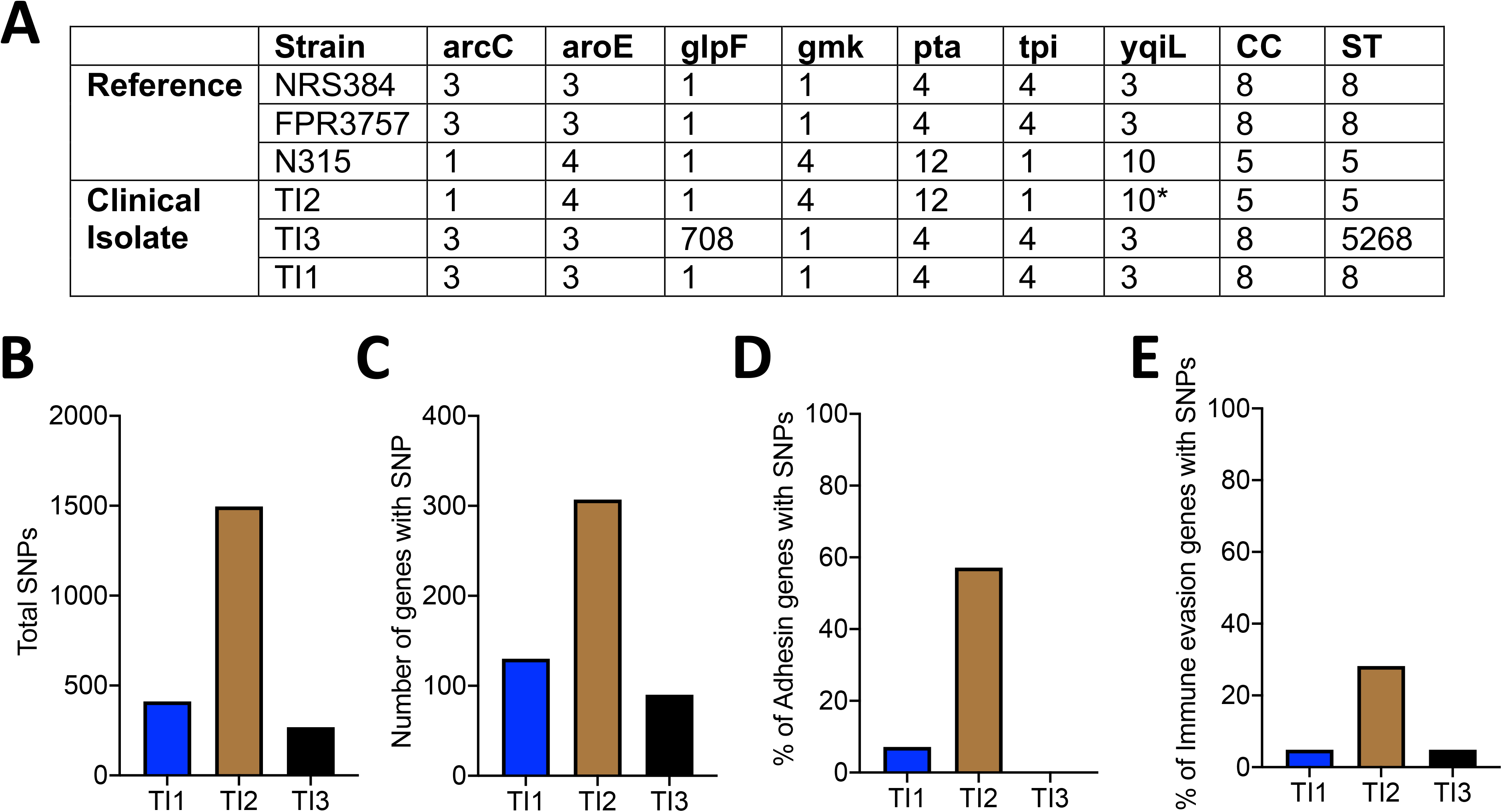
Genetic differences between clinical isolates of *S. aureus*. (A) Multilocus sequence typing (MLST) of the clinical isolates (TI1, TI2, TI3) and the reference strains NRS384 (USA300), FPR3757 (USA300), and N315 (USA100) listed with the allele ID numbers (* denotes closest match, no exact allele ID identified) for each MLST locus, clonal complex (CC), and strain type (ST). Enumerated single nucleotide polymorphisms (SNPs) as compared to reference strain (B), total number of genes with at least one SNP (C), and percentage of adhesin genes (D) or immune evasion genes (E), as designated in Fig. S8, with at least one SNP.

## Discussion

Using multiple clinical isolates of *S. aureus* from children with osteomyelitis, as well as two community-acquired strains from adult patients (NRS384 and LAC, originally isolated from skin (32, 33), we achieved reproducible development of HOM in mice. By generating stably bioluminescent bacteria, we were able to track infection development and location across time. Important for their utility in the study of HOM, the strains and clinical isolates tested produced frequent hind limb infections that did not rely upon direct bone inoculation or prior bone manipulation. Furthermore, mice did not become overtly septic immediately following injection and mostly survived the duration of the experiment, except for occasional hind-limb lameness or paralysis necessitating euthanasia after several weeks. The ease of tracking, early onset of infection, associated bone loss, and formation of characteristic abscesses all stand to make this a very effective model for the further study of HOM pathogenesis. Because BLI signal in bone correlated well with bacterial load, both *in vivo* and *in vitro*, destructive CFU assays are not required to assess effects of manipulations of either host or pathogen on bacterial burden in this model, and the bacteria can be tracked longitudinally in each mouse. Additionally, our HOM model mimics many of the hallmarks of juvenile HOM. Clinically, the most frequent site of infection is the metaphysis of the femur or tibia (12), a trend recapitulated in our studies. We also found several examples of abscesses located within the diaphysis of the femur and tibia, although this was not in the majority of samples. We observed rapid trabecular bone destruction, as early as 2-weeks post injection. However, significant cortical bone changes were rare, with only a few examples of periosteal reactive bone formation, along with one sequestrum. These infections persisted over the duration of our 5-week follow-up, but longer periods of study are warranted to fully characterize the chronic effects of these infections, as persistence and recurrence are some of the most challenging aspects of treating HOM patients.

Both BLI and CFU assays support the primary sites of infection after intravenous inoculation of all strains examined to be bone/joint and kidney/bladder. However, CFU assays also demonstrate low, but detectable, levels of live bacteria in femurs without distinct foci of bioluminescence or histologic changes such as abscess formation. Interestingly, in mice inoculated with the pediatric isolates, these BLI- femurs had trabecular bone volumes that were higher than the BLI+ femurs, but lower than those of control uninfected mice. The lack of bone volume differences between BLI+ and BLI- femurs in NRS384 and LAC infected mice may indicate a limitation of BLI in predicting bone loss, but this could also indicate inherent differences in the ability of certain strains to cause bone loss. Ultimately, more detailed assessments, including examination of other skeletal sites and imaging of bacteria in situ at high resolution, are needed to determine whether bone loss at BLI-negative sites is driven by systemic inflammation, direct effects of sparsely distributed bacteria in bone, or a combination of both.

Interestingly, by using multiple clinical isolates of *S. aureus*, we found overt differences in their pathogenicity ranging from infection incidence to predilection towards specific infection types (osteomyelitis versus septic arthritis). There were also more subtle differences between the isolates, such as the intensity of BLI signal in the BLI+ legs and the lesser drop in bone volume of IT2-injected mice compared to the other two isolates, although this could be a consequence of the lower percentage of bone-centered infections, as we were only able to use microCT to analyze two BLI+ legs. LAC also produced relatively few HOM infections, but those that occurred tended to involve the tibia rather than femur. However, only TI2, and not LAC, displayed lower osteoblast cytotoxicity and intra-osteoclast proliferation. These findings coincide with previous work establishing that *S. aureus* clinical isolates differ in their virulence, pathogenicity, and toxicity, and that a lower cytotoxicity correlates with increased persistence *in vivo* (34, 35). We found major differences in specific tissue tropism among isolates. TI2 displayed a predilection towards promoting SA compared to the other strains, especially TI3, while LAC predominantly targeted the tibia over the femur. However, our knowledge of the clinical behavior of these isolates is limited to only one patient per isolate, so it is currently unclear how these findings translate to a broader population. Further work is needed to determine specific bacterial factors that may promote differences in infection incidence and tropism, as well as if the lower cytotoxicity of TI2 contributes to chronicity as seen with other isolates (34, 35). It will be interesting to see if the incidence of sequestra or periosteal reactions – characteristic changes of chronic osteomyelitis - increases over time past the 5 weeks we examined here.

Using genomic sequencing of the new clinical isolates, we have begun to explore how genomic content shapes the observed clinical differences. When the isolates were sequence typed, the TI2 isolate was shown to be of the USA100 clonal lineage whereas the other isolates were of the USA300 lineage. USA100 has a higher prevalence in hospital-acquired infections, while USA300 predominates community-acquired infections (36, 37). While none of the patients from which these isolates were taken had hospital-acquired infections, a family member of the TI2 isolate patient was reported to be a healthcare worker. Although we do not yet know the traits of TI2 that underlie its pathogenic characteristics, this study highlights the importance of examining other lineage strains besides USA300 as they are still circulating, are able to cause OM and SA, and could have differences in their pathogenicity. Sequencing these isolates is the first step in understanding these differences and gives us a resource to identify other genetically related strains. TI2 was found to have a number of polymorphisms in specific genes related to adhesins and immune evasion. This could explain the lower infection incidence as both surface adhesion and the ability to evade immune destruction are critical steps in infection initiation and propagation. Also, differences in adhesin expression may help explain the predilection of TI2 to promote SA, as opposed to OM. Despite the different clades, TI2 and LAC have similar overall infection incidences in our model. However, LAC displayed similar osteoblast cytotoxicity and intra-osteoclast proliferation as the other USA300 strains, whereas strain TI2 was significantly less cytotoxic. These results, taken together with infection incidence differences, suggest that the strain of bacteria could be important to note when comparing results from different models of osteomyelitis.

It is apparent in our study that many mice in our model develop kidney infections as well as OM in long bones, so it is difficult to determine with certainty whether OM is primary or secondary. However, there are mice that only develop infections in the tibia or femur, suggesting this to be the primary infection location, and we have not seen any examples of mice that develop OM after first having a kidney/bladder infection identified by BLI. Another limitation is that our study has focused only on MRSA isolates. Methicillin-susceptible strains of *S. aureus* are also a significant cause of hospital- and community-acquired infections that will require investigation.

Ultimately, our work demonstrates that HOM can be reproducibly initiated in immunocompetent mice using clinical isolates, and that utilizing multiple isolates is likely necessary to capture the range of human disease. This panel of bioluminescent strains will be critical for further studies to elucidate biological mechanisms underpinning pediatric HOM development and propagation, eventually aiding in more efficacious clinical treatments and therapies.

## Materials and Methods

### Animals

For these studies, we used 6-week old male C57BL/6 mice, with a mutation in the tyrosinase gene (B-6 albino; B6(Cg)-Tyr^c-2J^/J; Lane, 1973; JAX stock #000058, The Jackson Laboratory, Bar Harbor, ME (38)) which renders their fur albino and allows for improved bioluminescent imaging. For hematogenous osteomyelitis modeling, 2.5×10^6^ bacteria were injected in 100µL of PBS via the tail vein. Control, uninfected mice used for microCT analysis were age and sex matched. Mice were housed 5 per cage with food and water ad libitum and closely monitored throughout experiments for signs of morbidity until humanely euthanized by asphyxiation with CO_2_.

### Clinical isolates

Clinical isolates were obtained from the St. Louis Children’s Hospital and Barnes Jewish Hospital clinical microbiology laboratories by Stephanie Fritz and transferred to Deborah Veis in accordance Washington University Institutional Review Board approval (IRB 201805091). None of the patients required pediatric intensive care unit admission or intubation. Antibiotic susceptibility profiles are as follows:

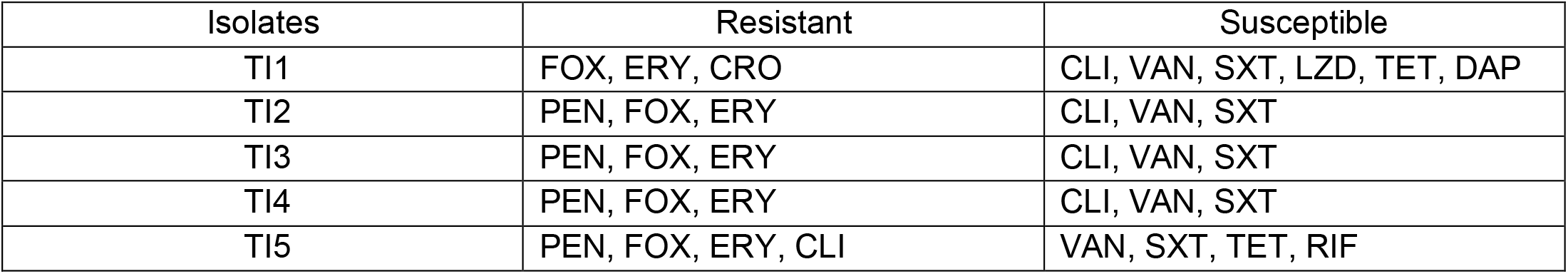
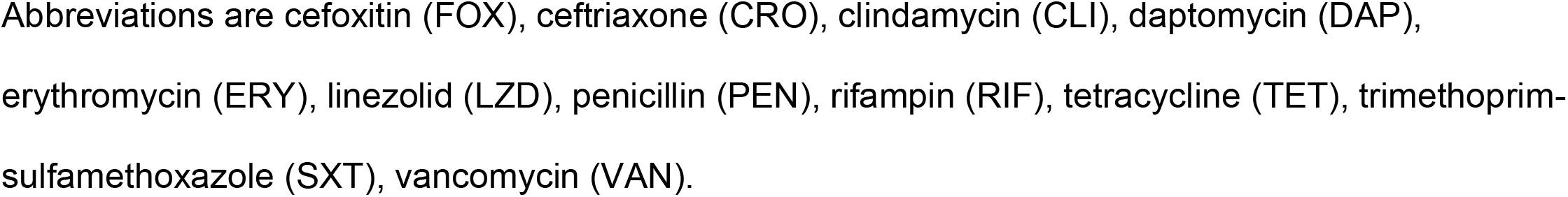

| Isolates | Resistant | Susceptible |
| --- | --- | --- |
| TI1 | FOX, ERY, CRO | CLI, VAN, SXT, LZD, TET, DAP |
| TI2 | PEN, FOX, ERY | CLI, VAN, SXT |
| TI3 | PEN, FOX, ERY | CLI, VAN, SXT |
| TI4 | PEN, FOX, ERY | CLI, VAN, SXT |
| TI5 | PEN, FOX, ERY, CLI | VAN, SXT, TET, RIF |

Abbreviations are cefoxitin (FOX), ceftriaxone (CRO), clindamycin (CLI), daptomycin (DAP), erythromycin (ERY), linezolid (LZD), penicillin (PEN), rifampin (RIF), tetracycline (TET), trimethoprim-sulfamethoxazole (SXT), vancomycin (VAN).

### Bacterial growth and preparation of inocula

All strains of *S. aureus* were routinely grown on tryptic soy agar (TSA) or in tryptic soy broth (TSB) (Fisher Scientific; Hampton, NH) at 1:200 shaking at 220 rpm overnight and subcultured 1:100 for 2 hours before injection. Bacteria were subsequently pelleted and resuspended in PBS to a final optical density at 600nm (OD_600_) of 1. Injections were prepared by further dilution of bacteria in PBS to a final count of 2.5×10^6^ in 100µL of PBS.

The NRS384 *S. aureus* strain used was already stably bioluminescent (27). The LAC strain of *S. aureus* used was AH1263, which was created by curing USA300-0114 of its Erm resistance (28). In order to produce stable bioluminescence in the clinical isolates, electrocompetent cells from each isolate were prepared and electroporated with the pRP1195 plasmid, as previously described (27). Clones containing the plasmid were selected for by chloramphenicol resistance and bioluminescence. Plasmid integration was then driven by a temperature shift to 43°C for 6-7 hours, as previously described (27), after which time bacteria were serial diluted onto TSA plates containing 5µg/mL and grown at 43°C overnight. Clones with bioluminescence were again selected, and 20% glycerol stocks made.

### Planktonic Growth Assay

Bacteria were streaked onto TSA plates and grown at 37°C overnight. Single colonies were selected and cultured in TSB shaking at 220 rpm overnight at 37°C. The overnight culture was then diluted in TSB 1:100 in fresh TSB. The OD_600_ was measured every 30 minutes for 5 hours.

### Bioluminescent imaging

*In vivo* bioluminescence imaging was performed on an IVIS 50 (PerkinElmer, Waltham, MA; Living Image 4.3.1) using 1-minute exposure, bin 8, FOV12cm, f/stop1, open filter. Mice were imaged under isoflurane anesthesia (2% vaporized in O_2_). Bacterial cultures grown *in vitro* were imaged using the same IVIS 50 using 1- second exposure, bin 8, FOV12cm, f/stop1, open filter. Total photon flux (photons/sec) was measured from fixed regions of interest (ROIs) set over the hindlimbs, as depicted (Fig. S2, or individual wells using Living Image 2.6.

### Micro-computed tomography (microCT) analysis

Hindlimb bones were fixed for 48 hours in 10% neutral buffered formalin, then stored in 70% ethanol. Bones were scanned *ex vivo* using a µCT40 (Scanco Medical, Switzerland) (10µm, 55 kVp, 145 µA, 8W, 300ms integration time) and analyzed with µCT Evaluation Program V6.6. For trabecular bone analyses, regions of interest were defined as 100 slices below the femoral growth plate. Outcome variables are reported in accordance with published consensus guidelines (39). Mice sacrificed earlier than desired end timepoint were not included in microCT analysis.

### Histology

After microCT analysis was performed, bones were decalcified for 14 days in 14% EDTA free acid, then stored in 70% ethanol. For tissue processing and staining, samples were submitted to the Washington University Musculoskeletal Histology and Morphometry Core for hematoxylin and eosin (H&E) or tartrate-resistant acid phosphatase (TRAP) staining. 2 to 3 sections were examined per mouse. Images were taken using an Olympus BX-51 microscope equipped with an Olympus DP27 camera and cellSens Standard software.

### Enumeration of colony forming units

At 3 weeks post-injection, tissues were harvested using a Bullet-Blender and NAVY lysis tubes (Next Advance, Inc., Averill Park, NY) at 4°C. To enumerate bacterial CFUs, the whole organ (tibia, femur, heart, kidney, or liver) was homogenized in TSB, then serially diluted in TSB, and plated on TSA plates at 37°C overnight for bacterial colony enumeration.

### Intracellular proliferation assay

Osteoclasts (OCs) generated from enriched bone marrow macrophages (BMMs) harvested from the long bones of 10- to 12- week-old wild type C57BL/6 mice and cultured in alpha-MEM with 10% FBS and 1:10 CMG 14-12 cell supernatant (containing the equivalent of 100 ng/mL of M-CSF) for 4 days to expand BMMs. BMMs were seeded into 12-well tissue-culture treated plates at 1.5×10^5^ per well and differentiated into OCs for 3 days in alpha-MEM with 10% FBS, 1:50 CMG 14-12 cell supernatant (20 ng/ml M-CSF), and 60 ng/ml GST-RANKL.

CFUs were enumerated after gentamicin protection assay, as previously described (29). Briefly, OCs were infected with *S. aureus* strains at an MOI 10:1 for 30 minutes. Cells were then washed with PBS and cultured in media containing antibiotic (0.3 mg/ml gentamicin) for 1 hour to kill extracellular bacteria. Then cells were again washed in PBS, and the 1.5-hour time point cells were lysed in sterile, cold molecular-grade H_2_O. 18-hour time point cells were replenished with media with M-CSF an RANKL until hypotonic lysis, as described above. Lysates were 10-fold serially diluted and plated on TSA plates for overnight incubation at 37°C.

### *S. aureus* culture supernatant concentration

Three colonies from each *S. aureus* strain were inoculated into 50 mL of RPMI supplemented with 1% casamino acids in a 250 mL Erlenmeyer flask. Flasks were stoppered and incubated at 37°C with shaking at 180 rpm for 15 hours. Bacterial cultures were centrifuged at 4,000 *× g* for 10 minutes at 4C, and culture supernatants were removed and filter-sterilized using a 0.22 µm filter. The supernatants were concentrated with an Amicon Ultra 3 kDa nominal molecular weight limit centrifugal filter unit (Millipore), according to manufacturer’s instructions. Concentrated supernatants were filter-sterilized again with a 0.22 µm filter, and aliquots of each supernatant were stored at −80°C until use in the cytotoxicity assay.

### Cytotoxicity assay

MC3T3-E1 cells obtained from the American Type Culture Collection (ATCC) were propagated according to ATCC recommendations. Cells were grown at 37°C and 5% CO_2_ with replacement of media every 2 to 3 days. For cytotoxicity assays, MC3T3 cells were seeded in 96-well tissue culture plates at a density of 5,000 cells in 200 µL media per well. After 24 hours, the media was removed and fresh media containing either 5% or 10% concentrated *S. aureus* culture supernatant or RPMI was added to the cell monolayers. The MC3T3 cells were incubated for an additional 22 hours, then cell viability was assayed using CellTiter 96 AQ_ueous_ One Solution Cell Proliferation Assay (Promega), according to manufacturer’s instructions. Survival was calculated for MC3T3 cells exposed to each supernatant condition as a percentage of survival relative to the MC3T3 cells exposed to media containing RPMI.

### *S. aureus* clinical isolate sequencing

DNA was isolated from overnight bacteria cultures using the DNeasy Blood and Tissue Kit (Qiagen Cat No 69504) per manufacturer’s instructions. Whole genome sequencing was performed at the Genome Technology Access Center at Washington University in St. Louis School of Medicine using Illumina NovaSeq 6000 with S4 flowcell and 2×150 paired-end reads. Sequence reads were aligned to the reference genome using NovoAlign. USA300_FPR3757 (NCBI accession number NC_007793.1) was used for the USA300 reference genome, and N315 (NCBI accession number BA000018.3) was used for the USA100 reference genome. The alignments were sorted with SAMtools. Picard Tools was used to remove duplicate reads. Variant files and BAM alignments for each clinical isolate were analyzed using Geneious Prime v. 2021.0.1. Submission of genomic sequencing data is in process at NCBI Sequence Read Archive (SRA).

### Statistical analysis

All data are represented as means with standard deviation. All data was analyzed by one-way analysis of variance (ANOVA) with Tukey’s multiple comparison *posthoc* test (GraphPad InStat), except for correlation data where the Pearson correlation coefficient was calculated, and Fig. 4A where differences between BLI- and BLI+ within each isolate was calculated by Student’s t-test (GraphPad InStat). P<0.05 was taken as significant.

### Ethics statement

All animal procedures were approved by the Institutional Animal Care and Use Committee of Washington University (IACUC protocol 20170025 and 19-1059), according to established federal and state policies outlined in the Animal Welfare Act (AWA) and enforced by the US Department of Agriculture (USDA), Animal and Plant Health Inspection Service (APHIS), USDA Animal Care. Clinical isolates were obtained under Washington University Institutional Review Board approval (IRB 201805091).

## Acknowledgements

This work was supported by National Institutes of Health grants R21 AR073507 and R01 AR070030 (D.J.V.) and by the Shriners’ Hospitals for Children grant 85117 (D.J.V.). P.M.R. was supported by a Skeletal Disorders Training Program T32 AR060719. J.E.C. was supported by R01AI132560 (NIAID), R01AI145992 (NIAID), and a Career Award for Medical Scientists from the Burroughs Wellcome Fund. K.R.E. was supported by the Childhood Infection Research Program T32 AI095303. C.A.F. was supported through T32GM007347 (NIGMS) and is supported by F30AI138424 (NIAID). The bioluminescent imaging was performed at the Washington University School of Medicine Molecular Imaging Center, supported by NIH P50 CA094056 (Molecular Imaging Center) and NCI P30 CA091842 (Siteman Cancer Center Small Animal Cancer Imaging Shared Resource). The Washington University Musculoskeletal Research Center, supported by P30 R074992, provided resources for microCT through its Structure & Strength Core, and histology via its Histology & Morphometry Core. Sequencing was performed by the Washington University Genome Technology Access Center.

We thank Crystal Idleburg and Samantha Coleman for expert histology, Roger Plaut for providing the lux operon plasmid pRP1195, and Julie Prior and Katie Duncan for their bioluminescence imaging assistance.

---

**Figure S1.**
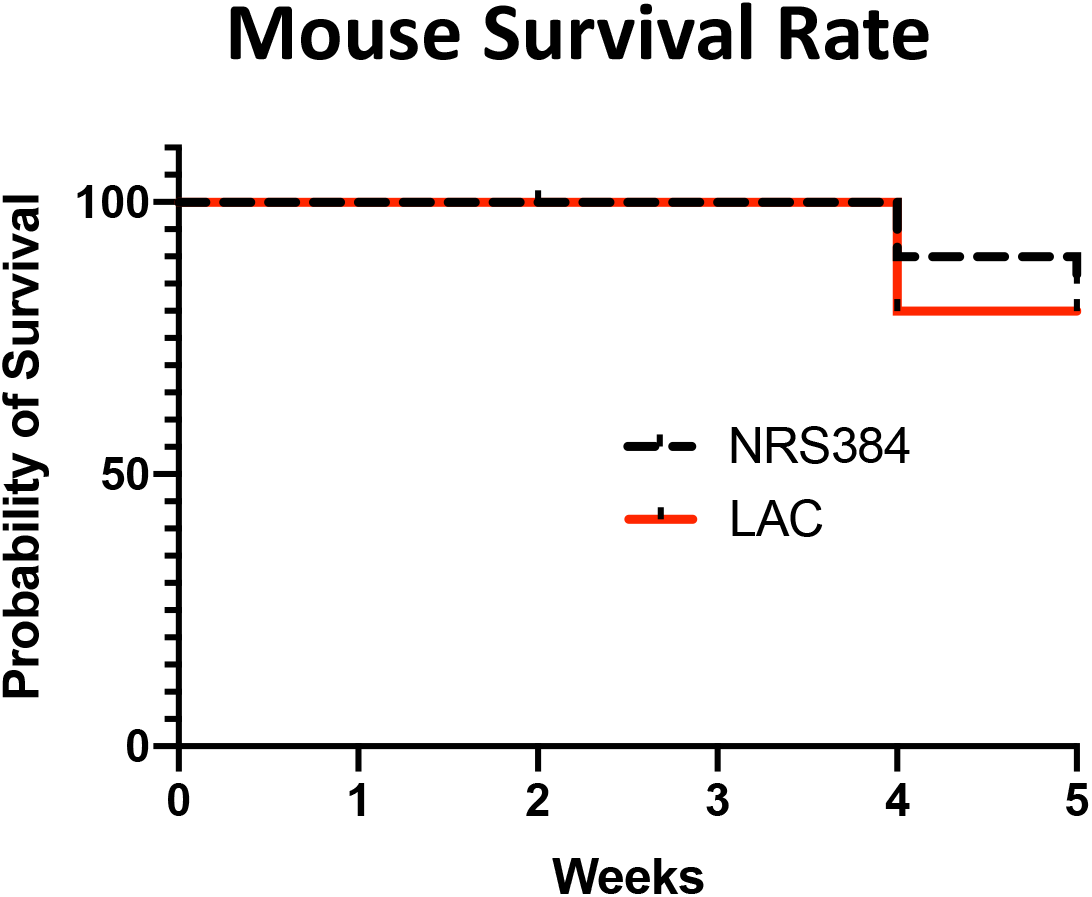
Percentage of mouse survival by week after injection with either NRS384 or LAC. Mice were sacrificed early (3 or 4 weeks post-injection) if lameness or hind-limb paralysis was observed.

**Figure S2.**
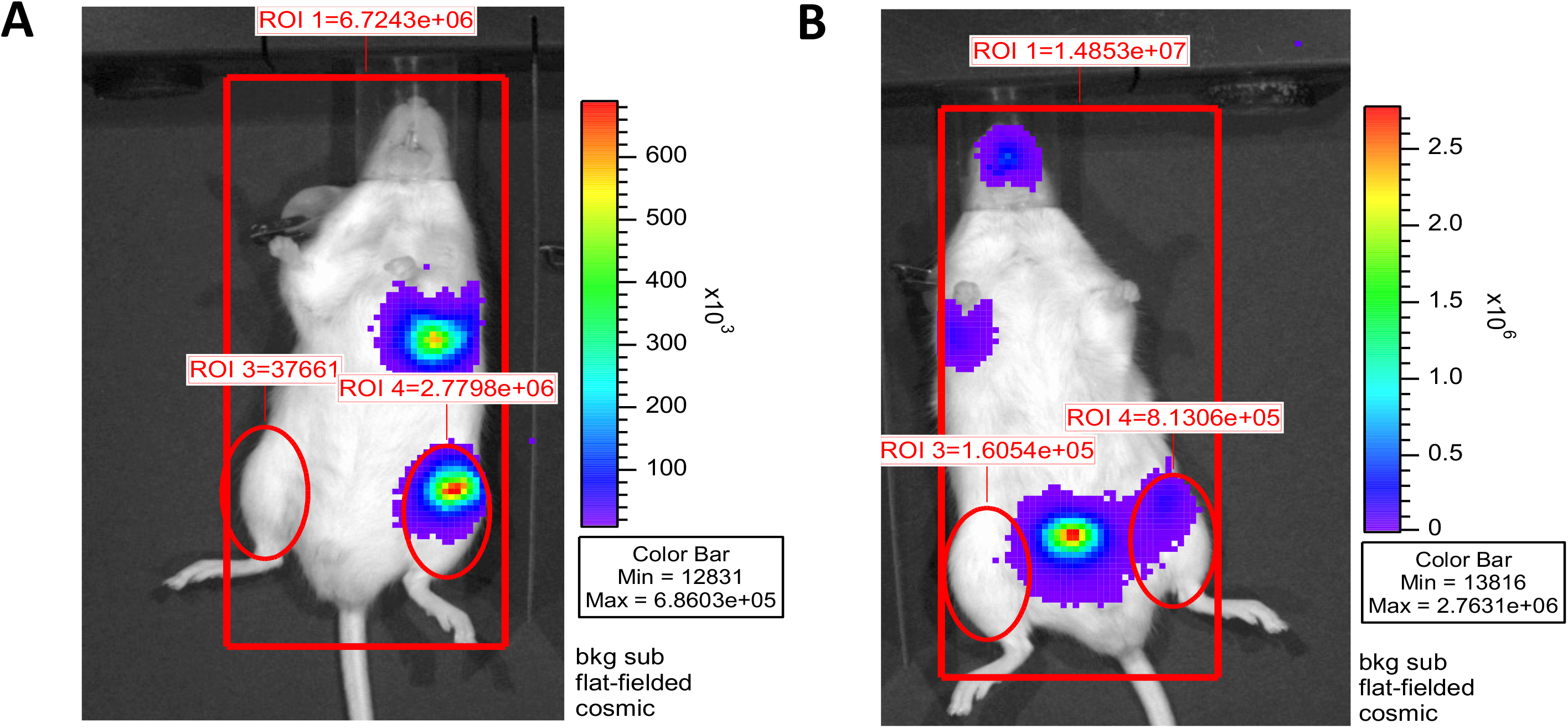
Explanation of BLI signal measurements. Regions of interest (ROIs) used to measure the BLI signal from each hind limb are shown. (A) An example of a mouse with one BLI+ (left) leg and one BLI- (right) leg. BLI+ designation relies upon a discrete BLI signal focus originating from the left leg exceeding a signal of 70,000 photons/second. (B) An example of a mouse with one BLI+ (left) leg and one BLI- (right) leg. The right leg of this mouse is designated as BLI- even though it has a BLI signal of 1.6054×10^5^ because there is no clear focus of signal originating from the leg and high BLI signal originating from the bladder at the edge of the leg ROI.

**Figure S3.**
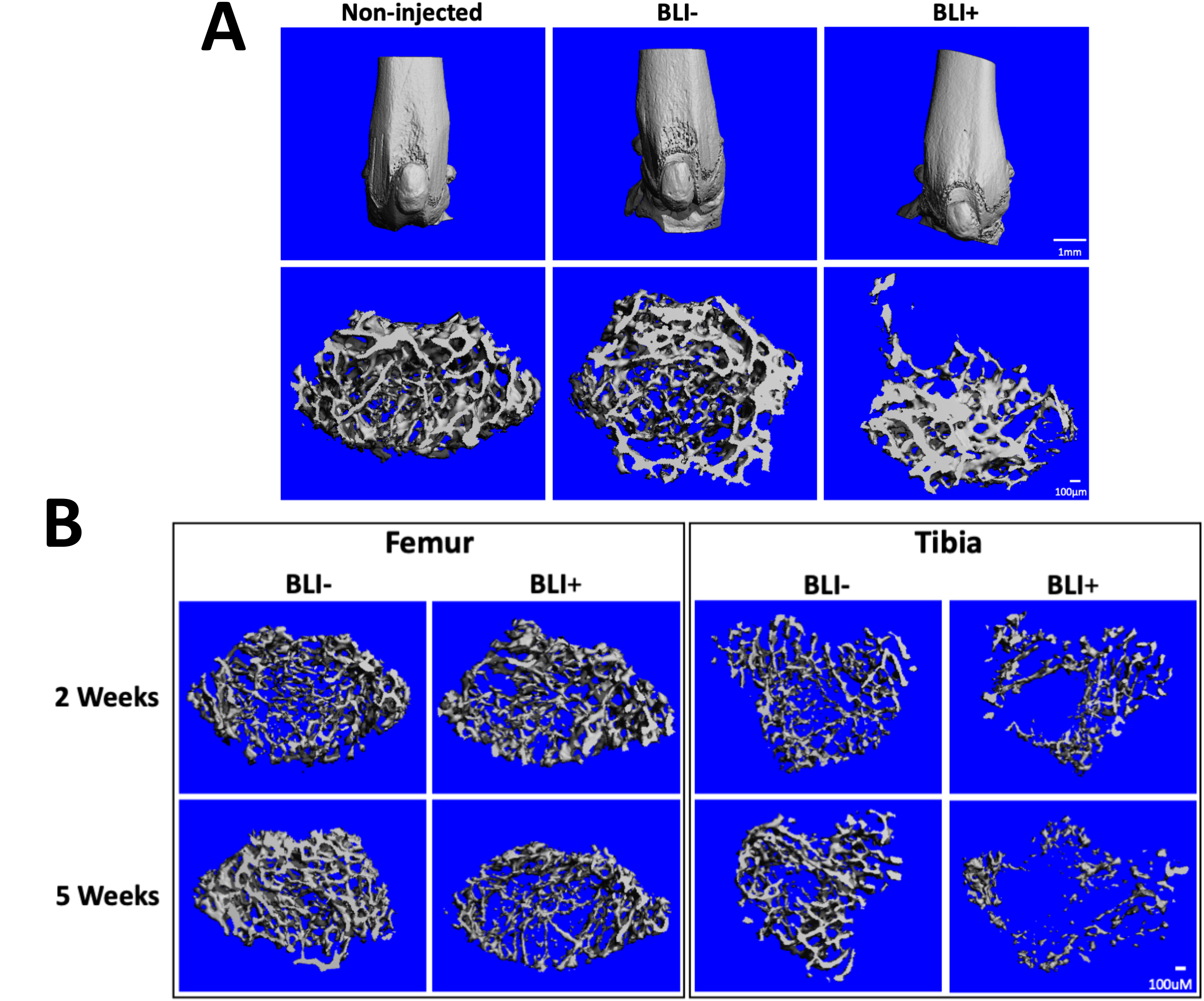
Effects of HOM on different bone compartments. (A) Representative microCT reconstructions of cortical (upper panels) and trabecular (lower panels) bone of femurs from control (Non-injected) mice or BLI- and BLI+ femurs after NRS384 injection. (B) Representative microCT reconstructions of trabecular bone of BLI- and BLI+ femurs and tibias from LAC injected mice measured at 2- or 5- weeks post injection. Scale bars 1 mm for cortical scans and 100 µm for trabecular scans.

**Figure S4.**
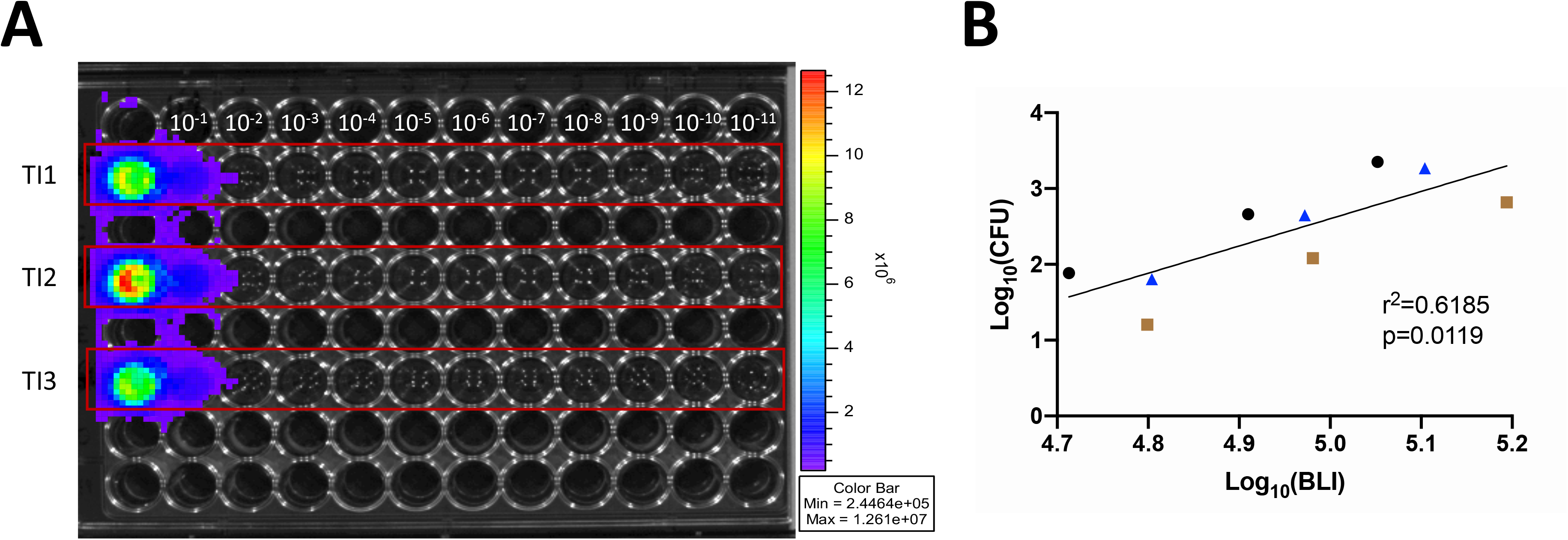
*In vitro* CFU and BLI correlation with *lux* operon in clinical isolates. (A) BLI image of serially diluted TI1, TI2, and TI3. Bacteria were grown overnight in TSB at 1:200 shaking at 220 rpm overnight and subcultured 1:100 for 2 hours before dilutions. (B) Correlation of the BLI signal from 10^-4^, 10^-5^, and 10^-6^ dilutions (A and B) from each isolate and the CFU count for these dilutions. CFUs enumerated from sampling of each dilution grown overnight on TSA. Blue triangle, TI1; brown squares, TI2; black dots, TI3.

**Figure S5.**
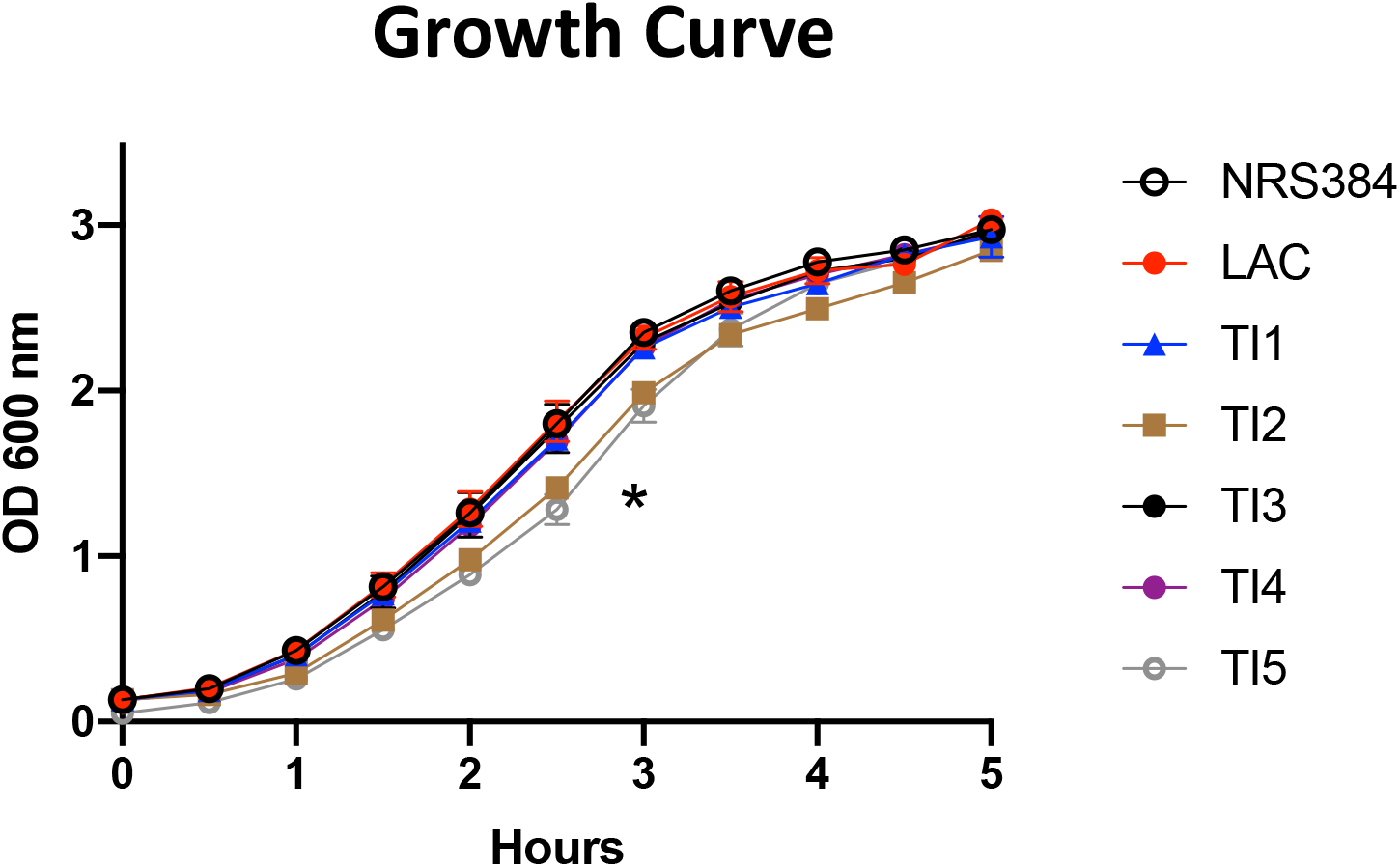
Bacterial growth assay. The OD_600_ of each strain and isolate was measured every 30 minutes for 5 hours to determine growth curves. p=0.0011, TI2 and TI5 compared to all other strains, by repeated measures two-way ANOVA with Geisser-Greenhouse correction and Tukey’s post-hoc test.

**Figure S6.**
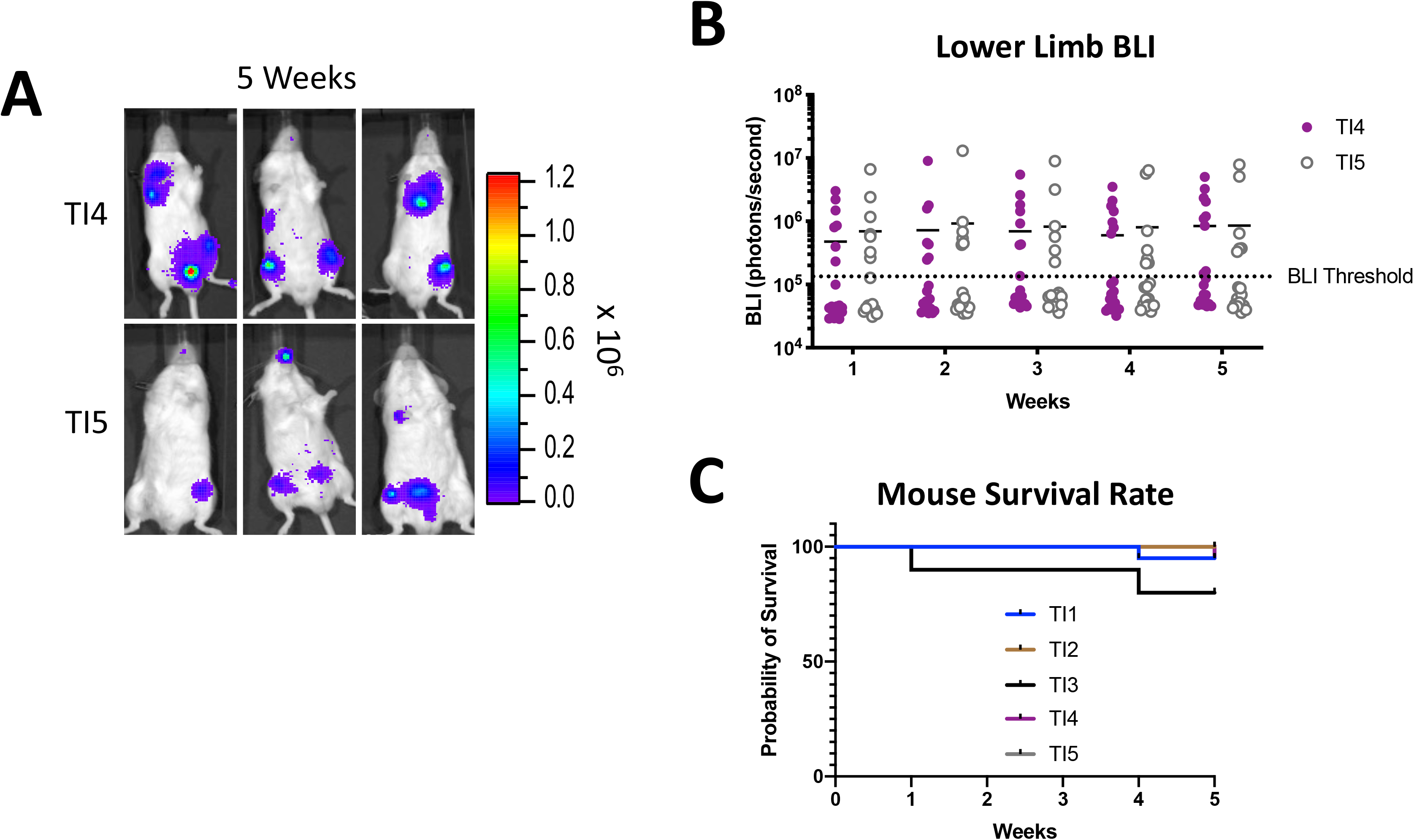
Characterization of additional clinical isolates of *S. aureus*. (A) *In vivo* bioluminescent imaging (BLI) of three representative mice for each isolate at 5 weeks post-injection. (B) The quantification of BLI signal measured in the hind limb of each injected mouse over the 5 weeks post-injection (n=20/strain; BLI Threshold 70,000 photons/second). (C) Percentage of mouse survival after injection with each clinical isolate. TI2, TI4, and TI5 injected mice all had 100% survival.

**Figure S7.**
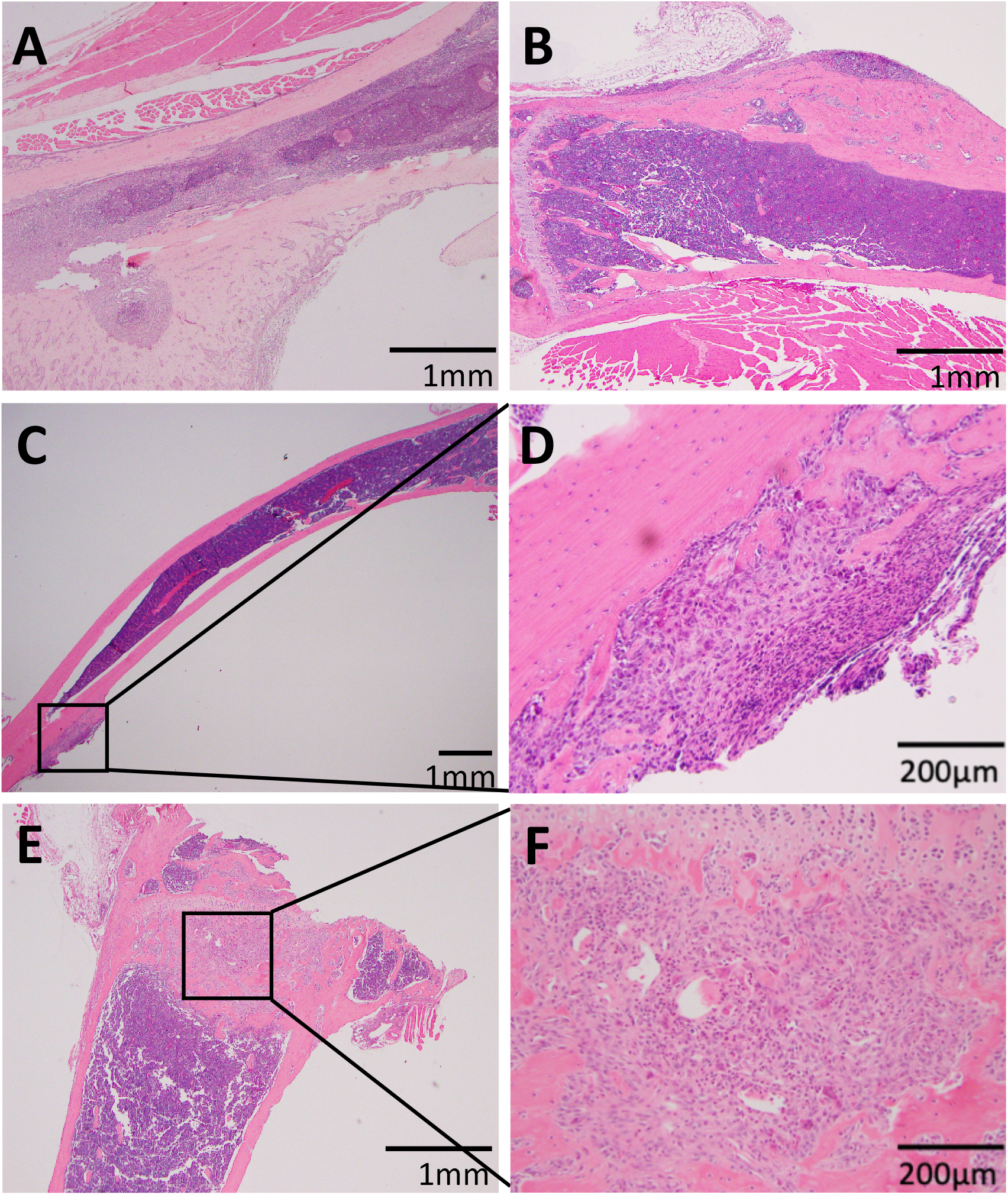
Unusual histological formations with HOM infections. (A) Abscess formed along the periosteum of the distal tibia with robust periosteal bone formation encompassing the abscess from a TI3-injected mouse. (B-D) Inflammation with reactive bone formation on the periosteal surface of two tibias of LAC-injected mice (D is higher magnification of black boxed area in C). (E,F) Fibrosis and new bone formation directly under the growth plate of a tibia from a TI2 injected mouse.

**Figure S8.**
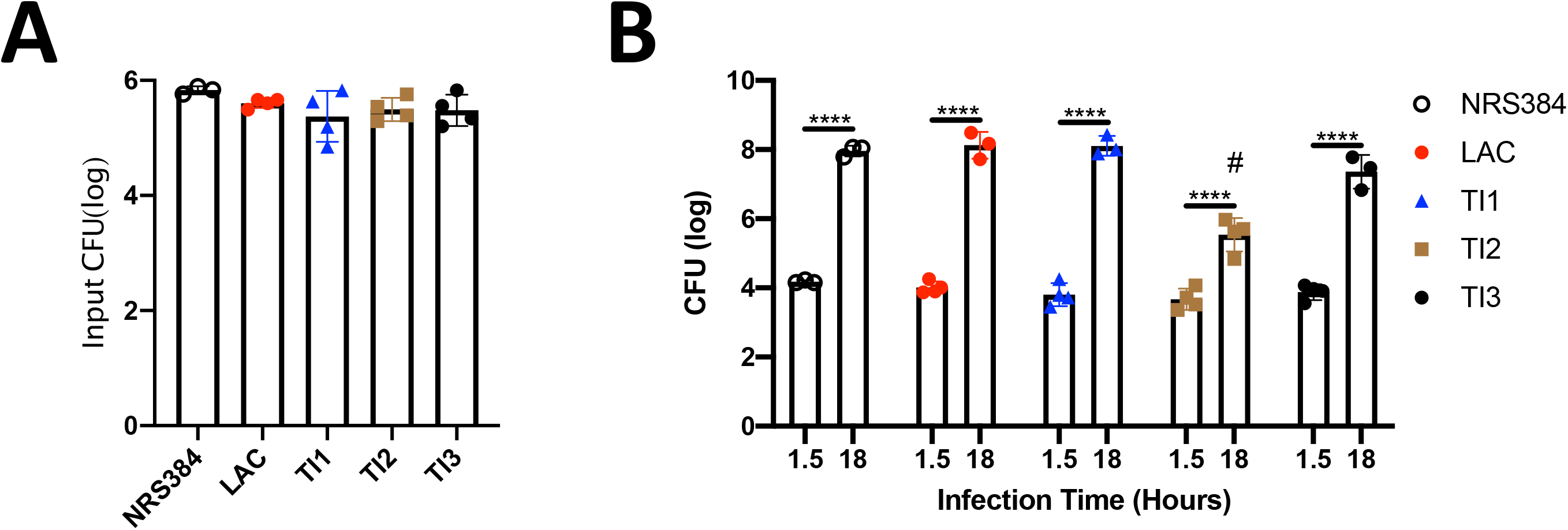
Osteoclast intracellular proliferation assay. (A) Confirmation of equivalent input CFU for each strain from the bacteria-containing media used to infect osteoclast cultures. (B) Intracellular CFU measured from lysed osteoclasts 1.5 or 18 hours post infection. Each dot represents a biological replicate (cells cultured from a different mouse on a different day). There is no difference in phagocytosis of bacteria (1.5 hour timepoint) in any strain. ****p<0.0001, 1.5 hour vs 18 hour timepoints, and #p≤0.0003, 18 hour TI2 vs all other groups, by Two-way ANOVA with Tukey’s post-hoc test.

**Figure S9.**
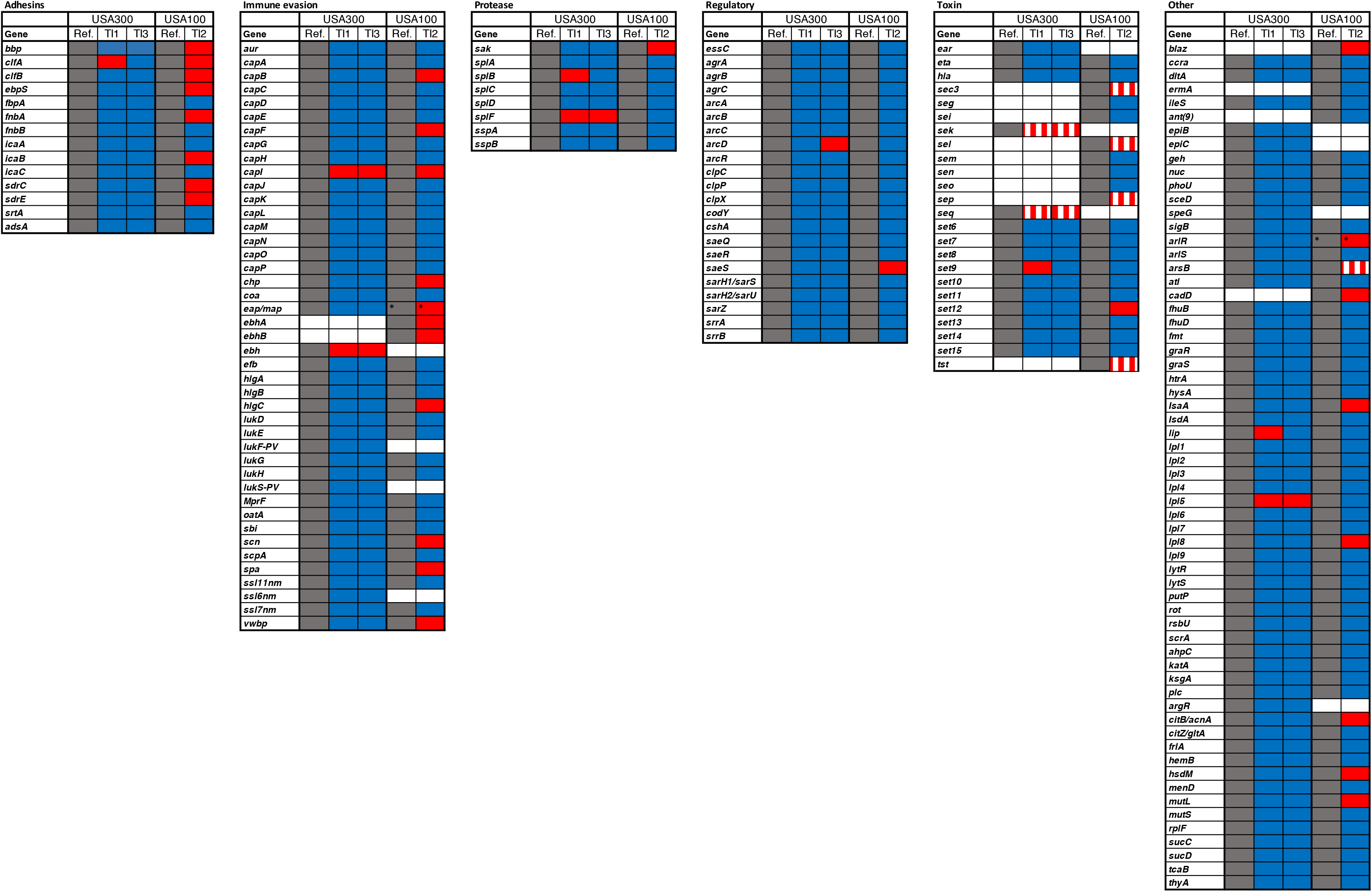
Clinical isolate polymorphisms by gene. 173 virulence associated genes (modified from (31)) and their presence, absence, and sequence relative to the reference strain. USA300_FPR3757 is the USA300 reference genome, and N315 is the USA100 reference genome. Grey = present in reference genome, White = absent in reference genome, Blue = present and identical sequence to reference genome, Red = present with at least one SNP, and Red/White stripped = absent in isolate but present in reference genome. * denotes a truncated gene. Genes are organized by function, which is listed above each table.

## Notes

### Competing Interest Statement

The authors have declared no competing interest.

